# Karyotype instability varies by species and genotype combination in allohexaploid *Brassica*

**DOI:** 10.1101/2023.09.22.559010

**Authors:** Daniela Quezada-Martinez, Jacqueline Batley, Annaliese S. Mason

## Abstract

Synthetic *Brassica* allohexaploids (2n = AABBCC) do not exist naturally but can be produced between six different parent species combinations, and can be used to investigate processes of polyploid formation and genome stabilization. In this study, we investigated hybridization potential, accumulation and frequency of copy number variants (CNVs), fertility, and karyotype stability in advanced generations of diverse allohexaploid genotypes belonging to different *Brassica* allohexaploid species combinations (NCJ types: *B. napus* × *B. carinata* × *B. juncea*; junleracea types: *B. juncea* × *B. oleracea*; naponigra types: *B. napus* × *B. nigra*; and carirapa types: *B. rapa* × *B. carinata*). Only 3.2% of allohexaploid plants investigated were euploids, with high frequencies of rearrangements. Significant differences between genotypes and between lineages within parent genotype combinations were found for frequencies of euploids and rearrangements, with one NCJ line showing relatively high karyotype stability. Hybridization between different allohexaploids was mostly achievable, with 0 - 4.6 seeds per flower bud on average, and with strong effects of maternal genotype. Novel hybrids between allohexaploid lineages showed similar fertility and stability to their parents. In the novel hybrid population, a significant correlation was observed between the inheritance of A-genome chromosome fragments (relative to C-genome fragments) and the total number of seeds produced per plant (*r* = 0.24). Our results suggest that synthetic *Brassica* allohexaploids can develop genomic stability, but that this occurs at very low frequencies, and may not always be under selective pressure due to the unpredictable relationship between fertility and genome composition in these hybrid types.

**Article Summary:** *Brassica* plants with three sets of chromosomes (allohexaploids) do not exist in nature, but can be made from combinations between six different species. Here, we compared different combinations to see which are the most genomically stable and fertile, and found major differences between all allohexaploid types as well as one putatively stable line. Crosses between allohexaploid types could also be achieved in most cases, although hybrids were not more stable or fertile than their parents.

## Introduction

Polyploidy is widely distributed in flowering plants, with recurrent whole genome duplication events dating back millions of years (Jiao et al. 2011). In general, there are two major polyploid types: autopolyploids (origin of additional chromosome sets from within the same species or even individual) and allopolyploids (two or more different chromosome sets from different species). Many crops are successful polyploids, such as wheat, oats, and rapeseed (Wendel 2000). Polyploidy can also have advantages such as increased cell size, growth rate, or organ size (Otto and Whitton 2000), fixed heterosis and gene redundancy (Comai 2005), and different soil and climate adaptations (Mousavizadeh et al. 2022). Unfortunately, many newly synthesized polyploids tend to be less fit, mostly due to genome instability in the context of meiosis (Pelé et al. 2018). In sexually reproducing plants, meiosis plays a fundamental role in ensuring correct chromosome recombination and inheritance into the next generations. In polyploids, the meiosis process becomes more convoluted since there are more possible pairing options per chromosome. Hence, the most common meiotic abnormality during meiosis in polyploids corresponds to chromosome associations involving multiple homologs or homoeologs (Grandont et al. 2013). These kinds of meiotic irregularities can lead to copy number variants (CNVs), aneuploid progeny, and non-viable gametes, making the stabilization of new polyploids a major challenge.

Studies trying to understand how polyploids might stabilize meiosis have been done in many species, and major loci and genes have been identified in Arabidopsis (e.g. (Yant et al. 2013; Seear et al. 2020)), wheat (Sears and Okamoto 1958; Griffiths et al. 2006; Bhullar et al. 2014; Rey et al. 2017), and *Brassica* (Jenczewski et al. 2003; Gaebelein et al. 2019b; Higgins et al. 2021), among others. In wheat polyploids, a major locus controlling homoeologous pairing has been identified as *Ph1* (*Pairing homoeologous 1*) located on chromosome 5B. In *B. napus* haploids (2n = AC), a major locus, *PrBn,* was identified as responsible for variation in pairing behavior (Jenczewski et al. 2003), although a similar effect of this locus was not observed in allotetraploid plants (Grandont et al. 2014). In a segregating *B. napus* population where homoeologous recombination events were quantified, three quantitative trait loci (QTL) were found on chromosomes A03 (23.3 - 26.3 Mb), A09 (11.1 - 23.9 Mb), and C07 (42.2 - 43.4 Mb) (Higgins et al. 2021). The QTL located on chromosome A09 accounted for up to 58% of the variability observed in homoeologous pairing (Higgins et al. 2021). When comparing gene expression using RNA-seq data collected from meiocytes, an ortholog of the meiotic gene *RPA1C* was found to be a candidate based on differential gene expression analysis (Higgins et al. 2021). In allohexaploid *Brassica* produced from the cross [(*B. napus* × *B. carinata*) × *B. juncea*], several QTL were identified on chromosomes A03, A04, A10, B05, B06, B08, and C03 that were putatively associated with meiotic behaviour (Gaebelein et al. 2019b).

Trigenomic allohexaploid *Brassica* (2n = 54 = AABBCC) do not exist in nature but can be produced via different cross combinations of the diploid and allotetraploid species (Gaebelein and Mason 2018). In the *Brassica* genus the diploid species *B. oleracea* (2n = 18 = CC)*, B. rapa* (2n = 20 = AA), and *B. nigra* (2n = 16 = BB) hybridized to form the allotetraploid species *B. napus* (2n = 38 = AACC)*, B. carinata* (2n = 34 = BBCC), and *B. juncea* (2n = 36 = AABB) (U 1935). The most common method to produce allohexaploids is the cross between *B. carinata* × *B. rapa* followed by induced chromosome doubling (e.g. via colchicine treatment), and this new synthetic type is known as “carirapa” (Tian et al. 2010; Zou et al. 2010; Gaebelein and Mason 2018). It is also possible to cross between *B. juncea* × *B. oleracea,* also followed by induced chromosome doubling (e.g. (Mwathi et al. 2019)), where the final hybrid is known as “junleracea”, or between *B. napus* × *B. nigra* followed by induced chromosome doubling to produce “naponigra” hexaploids. A fairly new method to produce allohexaploids relies on unreduced gamete production in a two-step hybridization method between [(*B. napus* × *B. carinata*) × *B. juncea*], referred to as “NCJ” types (Mason et al. 2012). Although trigenomic *Brassica* allohexaploids can be made via many combinations of species, the number of genotype combinations which have successfully produced a hybrid is very limited. To produce a trigenomic *Brassica* allohexaploid two or more different species are usually crossed and the success might depend on factors such as the direction of the cross, the ploidy of the parents, species origin (Morgan et al. 2021), temperature, and genetic variation (Bjerkan et al. 2020) on top of the pre- and post-fertilization barriers. Up until now, crosses between different allohexaploid types have not been produced or investigated. If crosses between different allohexaploid types can be achieved, this could help to increase genetic diversity in the limited existing material.

Synthetic allohexaploid *Brassica* are usually meiotically unstable to a greater or lesser degree, with strong genotype-specific effects (Gaebelein et al. 2019b). Meiotic instability resulting in non-homologous chromosome recombination and aneuploidy has been observed in the initial generations of each allohexaploid type in the majority of genotypes tested so far (Tian et al. 2010; Mason et al. 2014; Gupta et al. 2016; Zhou et al. 2016; Gaebelein et al. 2019a, b; Mwathi et al. 2020). To date, very high meiotic stability (as assessed by bivalent frequency) has only been observed in a carirapa allohexaploid derived from one specific *B. rapa* cultivar (Gupta et al. 2016). There is also evidence for relatively high stability (as assessed by generational frequency of 2n = 54 chromosome complements) in several other carirapa lines, although this was rare in a large set of characterized genotypes (Tian et al. 2010). There is also some evidence in different types of hexaploids for a generational increment in meiotic stability (Zhou et al. 2016). However, other studies suggest that aneuploid plants have a tendency to produce more aneuploid offspring (Iwasa 1964), and that existing non-homologous recombination events may also cause problems in subsequent generations (Mason et al. 2015; Mwathi et al. 2019). Despite previous work on this topic, it has to date been difficult to systematically compare genomic stability between allohexaploid combinations and genotypes, due to the wide range of different methodologies used to assess this.

In this study, we aimed to compare genome stability between eight genotypes (nine lineages) of different synthetic *Brassica* hexaploids resulting from three cross combinations type by assessing the frequency of novel CNVs arising in a single generation between sibling plants, in order to comprehensively determine differences between lines. We also assessed if karyotype changes were reproducible or random across the allohexaploid genomes after polyploidisation. We further aimed to determine how feasible it is to combine new *Brassica* allohexaploid types via the success of hand-crossing between and within four allohexaploid types and the fertility and karyotype stability of resulting progeny by assessing CNVs in the progeny.

## Materials and methods

### Plant material

Parental *Brassica* genotypes used to produce NCJ, junleracea, and naponigra allohexaploids were *B. napus* “Surpass400_024DH”, “Boomer”, “Ag-Spectrum”, “MSL Express”, and “Ningyou7”, from here on referred to as N1, N5, N6, N8, and N9 respectively; *B. carinata* “195923.3.2_01DH”, and “94024.2_02DH” (C1 and C2 respectively); and *B. juncea* genotypes “JN9-04”, “Purple leaf mustard”, and “B578” (J1, J2, and J3, respectively). *Brassica oleracea* genotype “TO1000” is referred to as O1, and *B. nigra* genotypes “Junius”, “IX7”, and “IX13”are referred to as I1, I2, and I3, respectively (Mason et al. 2010, 2012; Gaebelein et al. 2019a; Mwathi et al. 2020). Carirapa hexaploid parental genotypes used in the cross are as follows: *B. carinata* “CGN03943” × *B. rapa* “CGN03907” produced carirapa C05, *B. carinata* “CGN03949” × *B. rapa* “Ankangzhong” produced carirapa C13, and a cross between two different carirapa - C21 (*B. carinata* “CGN03983” × *B. rapa* “Wulitian YC”) and C28 (*B. carinata* “CGN03995” × *B. rapa* “Baijian 13”) - produced carirapa C2128 (Tian et al. 2010) .

### Genome stability analysis

The following *Brassica* parental genotypes were selected as controls: N1, N5, C1, C2, J1, J2, and O1. *Brassica* hexaploid genotypes were selected based on total seed number. From the NCJ type, one lineage from the N1C1.J1 genotype (generations H_5-6_), N1C2.J1 (generation H_6_), N6C2.J2 (generations H_5-6_) and two lineages from genotype N5C2.J1 (lineages described as a and b at the end of the genotype name, generations H_5_ and H_4_, respectively) were selected (Mason et al. 2012). From the carirapa type, one lineage per genotypes C05 (generation H_10_), C13 (generation H_8-9_), C2128 (generation H_6_) were chosen (Tian et al. 2010). From the junleracea type, one lineage of genotype J3O1 (Mwathi et al. 2020) was selected (H_4_ generation). From each *Brassica* control, five plants were grown (35 plants in total). In the case of the allohexaploids NCJ and carirapa, five different sibling lines per lineage were selected four to five plants per line were grown, giving a total of 24-25 per lineage, with a total of 124 plants from NCJ and 74 plants from carirapa. In the case of the junleracea type, four different sibling lines were selected and five plants were grown per sibling line, with a total of 20 plants (Supplementary Table 1).

### Crossings between four allohexaploid types

The following NCJ combinations were selected: N1C1.J1 (seven plants, H_3_ generation), N1C2.J1 (seven plants, H_4_ generation), N5C2.J2a (nine plants, H_3_ generation), N5C2.J2b (five plants, H_2_ generation), N6C2.J2 (nine plants, H_3_ generation), N4C2.J1 (eight plants, H_1_ generation), N5C2.J1 (three plants, H_2_ generation), and N7C1.J1 (five plants, H_2_ generation) (Mason et al. 2012). The *Brassica* carirapa allohexaploids selected were C05 (10 plants, H_8_ generation), C13 (seven plants, H_6_ generation), and C2128 (five plants, H_4_ generation) (Tian et al. 2010).

Cuttings of *Brassica* naponigra were used for crossing. The selected genotype combinations were N8.I1 (two cuttings), N8.I2 (two cuttings), N8.I3 (three cuttings), N5.I2 (four cuttings), and N9.I3 (four cuttings) (Gaebelein et al. 2019a). One genotype from *Brassica* junleracea was selected for crossing: J3O1 (five plants, H_2_ generation) (Mwathi et al. 2020).

### F_1_ hybrids and parents

*Brassica* F_1_ hybrids produced from crosses between different allohexaploid types were selected based on the number of seeds produced and the genotype combination. Allohexaploid genotypes selected and the individual parent plants that were used to produce the F_1_ hybrids were as follows: N1C1.J1 × N6C2.J2 (plant #2), N6C2.J2 (plant #3) × N5.I2, N6C2.J2 (plant #1) × C13 (plant #1), N6C2.J2 (plant #1)× J3O1 (plant #1), C13 (plant #2) × C05, C13 (plant #2) × J3O1 (plant #1), C13 (plant #3) × N8.I3, and N8.I3 × J3O1 (plant #2). Three plants were grown from each of the parent genotypes and F_1_ hybrid combinations for further analysis.

### Plant growth conditions, fertility, and plant data collection

Seeds were germinated in quick pot trays under greenhouse conditions at Justus-Liebig University (Giessen), then transferred into 1 L pots after reaching the two to three true-leaf stage. Plants used for the genome stability assay were grown only in quick pots until discarded. Plants were watered as required with weekly fertilization, and were grown under a photoperiod of 16 h day and 8 h night in a temperature-controlled chamber. Plant protection was applied to control pests and diseases as needed. Days to flowering was measured by counting from the day of sowing until the first flower opened (developmental stage BBCH60).

Pollen viability was estimated by using two flowers per plant using an Amphasys Z30 (Amphasys AG, Switzerland) with the F chip and AF7 buffer, following provideŕs recommendations (Supplementary Table 2). Pollen viability was also established by collecting two flowers with mature pollen and placing the pollen grain of each flower (6 anthers) on a glass microscope slide, staining it with acetocarmine 1% (1 g of carmine powder in 100 mL of 45% acetic acid) followed by visualizing under the microscope (Supplementary Table 2). For acetocarmine staining, 300 pollen grains were counted per flower, where round and red-stained pollen grains were considered alive and shrunken, yellow or unstained pollen grains were counted as dead. In both pollen viability estimates, the value was expressed as a percentage viability based on the relative numbers of live and dead pollen grains per sample. The average pollen viability was established based on two flowers counted for each plant.

The total number of self-pollinated seeds produced per plant for the parents and F_1_ hybrids was obtained by covering the entire plant with a microperforated bag (0.5 mm, Crispac-Beutel, Baumann Saatzuchtbedarf GmbH, Waldenburg, Germany) after the start of flowering, and keeping the bags on the plant until seed harvesting.

### Crossing success

Once at the flowering stage, the different genotypes were crossed in one direction (Supplementary Table 3). In total, we aimed to produced 126 different crosses with 100 flower buds per cross combination (12 000 flower buds in total) out of which 40 crosses were between NCJ and naponigra, 24 crosses between NCJ and carirapa, eight crosses between NCJ and junleracea, five crosses between junleracea and naponigra, 15 crosses between naponigra and carirapa, three crosses between junleracea and carirapa, 28 crosses between NCJ, and three crosses between carirapa genotypes. The female or male parent plant used in each of the crosses was selected based on phenotypic observations of pollen availability, number and size of flower buds, and overall fitness of the plant. The female parent plant was selected first, based on relative number of inflorescences and relative number of flower buds compared to the other genotype involved in the cross, where the plant with the highest numbers of both was selected as the female parent. In crosses where one parent plant or all plants of a specific genotype produced very little to no pollen, these were also selected as the female parent. The male parent was selected as a default in each combination after the female parent was chosen. Some plants produced both high numbers of flower buds and large amounts of mature pollen, and were used as both the female and male parent in different crosses.

Crosses were done by using forceps to open the flower buds from the female parent to remove the anthers and to expose the stigma. Then, mature anthers from the selected male parent were collected and gently rubbed over the female parent stigma. The cross was then labeled and covered with a microperforated plastic bag (0.5 mm, Crispac-Beutel, Baumann Saatzuchtbedarf GmbH, Waldenburg, Germany) until harvest to prevent contamination with external pollen donors. The female parent is the first genotype named in the corresponding cross combination.

In parallel, 1-10 branches per plant were covered with microperforated bags to produce self-pollinated seeds. After seeds in the crossing and self-pollination bags were finished ripening, total siliques developed, number of seeds per silique, viviparous seeds (germinated seeds inside the silique), and total seeds (normal seeds plus viviparous seeds) were counted. The ratio of crossing success was calculated by dividing the total number of seeds obtained by the total number of flower buds crossed. The number of siliques that developed after pollination (the number of instances when after pollination the silique elongated and putatively developed seeds, unlike undeveloped siliques that died and fell off the plant a few days after pollination) were also quantified.

### DNA extraction and SNP genotyping

After the plants reached the three to four true-leaves stage, approximately 100 mg of leaf material was collected and stored at -20°C until processing. DNA was extracted using the CTAB method (Doyle and Doyle 1990). The samples were then treated with RNAse (Carl Roth, Germany) according to manufacturer’s instructions. DNA was quantified using a Qubit fluorometer and dsDNA BR assay kit (Thermofisher Scientific, Germany) following manufactureŕs instructions. DNA quality was also checked using agarose gel electrophoresis.

Lyophilized DNA samples were sent for genotyping using the Illumina Infinium *Brassica* 90K SNP array (Illumina, USA) following the manufactureŕs instructions. The initial filtering of the data and analysis was done as previously described (Quezada-Martinez et al. 2022). Briefly, in Genome Studio 2.0 (Illumina, USA) the analysis of the A and C genome SNPs was done using the recommended cluster file (Clarke et al. 2016) and automated clustering was used for the B genome. The top-hit (highest *e*- value) was determined for the A and C genome probes based on a BLAST to the *B. napus* Darmor-*bzh* v. 8.1 reference genome (Bayer et al. 2017). The positions used for the B genome probe were the ones provided in the Illumina Infinium 90K SNP array, based on an early version of the short-read *B. nigra* genome Ni100 (Perumal et al. 2020). The initial cleaning of the data involved the removal of non-specific SNPs and those SNPs where the genomic location was not able to be determined. SNPs with 100% no-calls across all individuals were also removed from further analysis. A total of 42 554 SNP markers were kept for further analysis, distributed across the A (12 577 SNPs), C (17 867 SNPs), and B (12 111 SNPs) subgenomes (Supplementary Table 4).

### Copy number variant analysis

To determine the presence or absence of a chromosome we used SNP data combined with Log_2_ R ratio values (see method as described in (Quezada-Martinez et al. 2022)). The approximate centromere locations for the A and C chromosomes were based on the *B. napus* Darmor-*bzh* v. 8.1 reference genome (Bayer et al. 2017) using estimates remapped from (Mason et al. 2016). For the B genome, the approximate centromeric locations were taken from the *B. nigra* Ni100 genome (Perumal et al. 2020).

Copy number variation was established based on experimentally-derived values as described in (Quezada-Martinez et al. 2022). Fixed events for duplications and deletions were classified as such if the event was observed for both homologs. If the rearrangement event was heterozygous, showing as present in only one homolog, the event was classified as segregating.

Initially, all copy number events were scored independently by chromosome and by plant. Secondarily, potential translocation events were assessed based on known homoeologous relationships between the subgenomes (Chalhoub et al. 2014; Perumal et al. 2020). For example, once a duplication event was observed, the corresponding homoeologous region was also looked at: if the top of A01 had a 2 Mb duplication, the primary homeologous region at the top of C01 was checked, and if a deletion or missing copy was present in this region the event was classified as a translocation. The final karyotype was drawn using the R package chromDraw (Janečka and Lysak 2016) in RStudio v.2022.07.1 and later modified using GIMP v.2.10.20.

### Statistical analysis

Multiple regression models were used to determine the effects of independent variables on the number of seeds produced per number of flower buds crossed (crossing success ratio). To account for heteroscedasticity in our model (right-skewed residuals, Breusch-Pagan test *p* = 0.005208), the final model was calculated using weights. Initially, a multiple regression model including all the variables was created (allohexaploid type combination, genotype of female parent, genotype of male parent, and pollen viability). After initial analysis of single term effects, pollen viability had no statistically significant effect (*p* = 0.06247) and it was removed from the final model. The new model incorporated the remaining two terms without interactions and it explained 61% of the variability observed in ratio of seeds produced.

For the remaining analysis, normality and homogeneity of the data was checked using the Shapiro-Wilk Test and Leveńs Test in R. If the data met both assumptions, it was followed up with a pairwise Student’s t-test. For the remaining data, if the data did not meet the criteria for normality and homogeneity then the non-parametric Kruskal-Wallis test was initially performed, followed by a post-hoc Dunńs test. P-values were adjusted using the Bonferroni method. Data correlations were done using the non-parametric Kendall correlation method. Categorical variables (e.g. subgenome) were compared using Pearson’s χ^2^ test. All statistical analysis was performed in R v.4.0.2.

## Results

### Unexpected genome instability in *Brassica* parental species

Five plants from each of seven different control genotypes from distinct established *Brassica* species (*B. napus* N1 and N5*; B. carinata* C1 and C2; *B. juncea* C1 and C2; *B. oleracea* O1) were analyzed for fixed (present in both homologous chromosomes) and segregating (present in one homologous chromosome) CNV events. Both events which were fixed per genotype across all five plants and events which were present only in one or a subset of plants per genotype were observed (Figure 1). Fixed genomic rearrangements were present in both *B. napus* genotype N1 and *B. carinata* C1 (Figure 1). Segregating events were also detected in genotypes *B. napus* N1 and N5, and *B. carinata* C1. Based on homoeology, duplication events observed in *B. napus* N1 are likely non-reciprocal translocation events between chromosomes A01/C01, A04/C04, C09/A10, and C09/A09. Three events involved an extra copy of the C genome replacing a region in the A genome, and two events involved an extra copy of the A genome replacing a region in the C genome. Loss of a whole chromosome was observed in three different plants (*B. napus* N1 and N5), affecting chromosome number (Figure 1, Figure 2).

**Figure 1.**
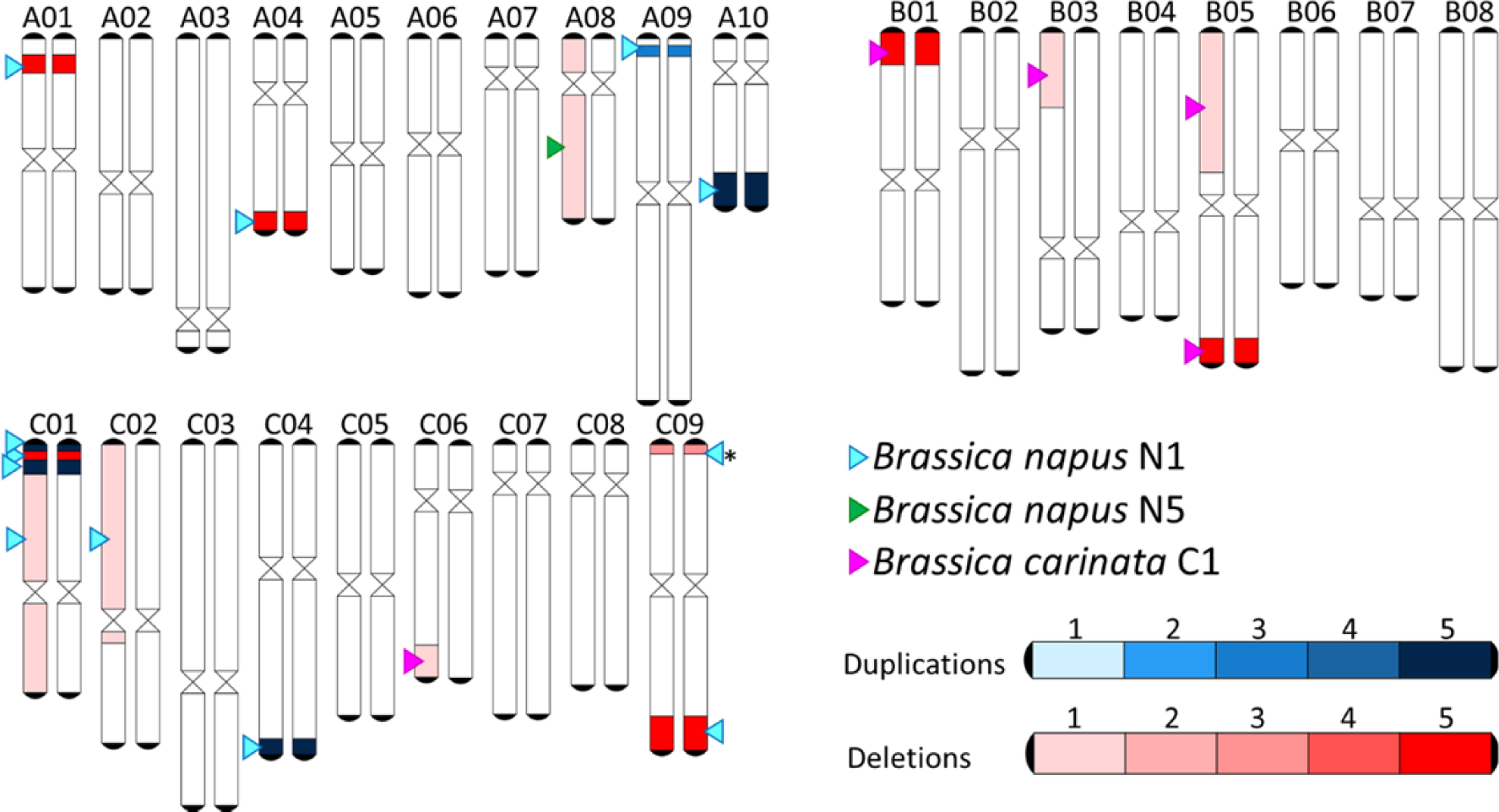
Combined molecular karyotype of *Brassica* species genotypes. Fixed (present in both homologs) and segregating events (present in one homolog) are represented. Frequency (in five assessed plants per genotype) is depicted in the legend. The genotype in which the rearrangement is present is indicated with an arrow head. *Deletion present as fixed in three plants and segregating in two plants.

**Figure 2.**
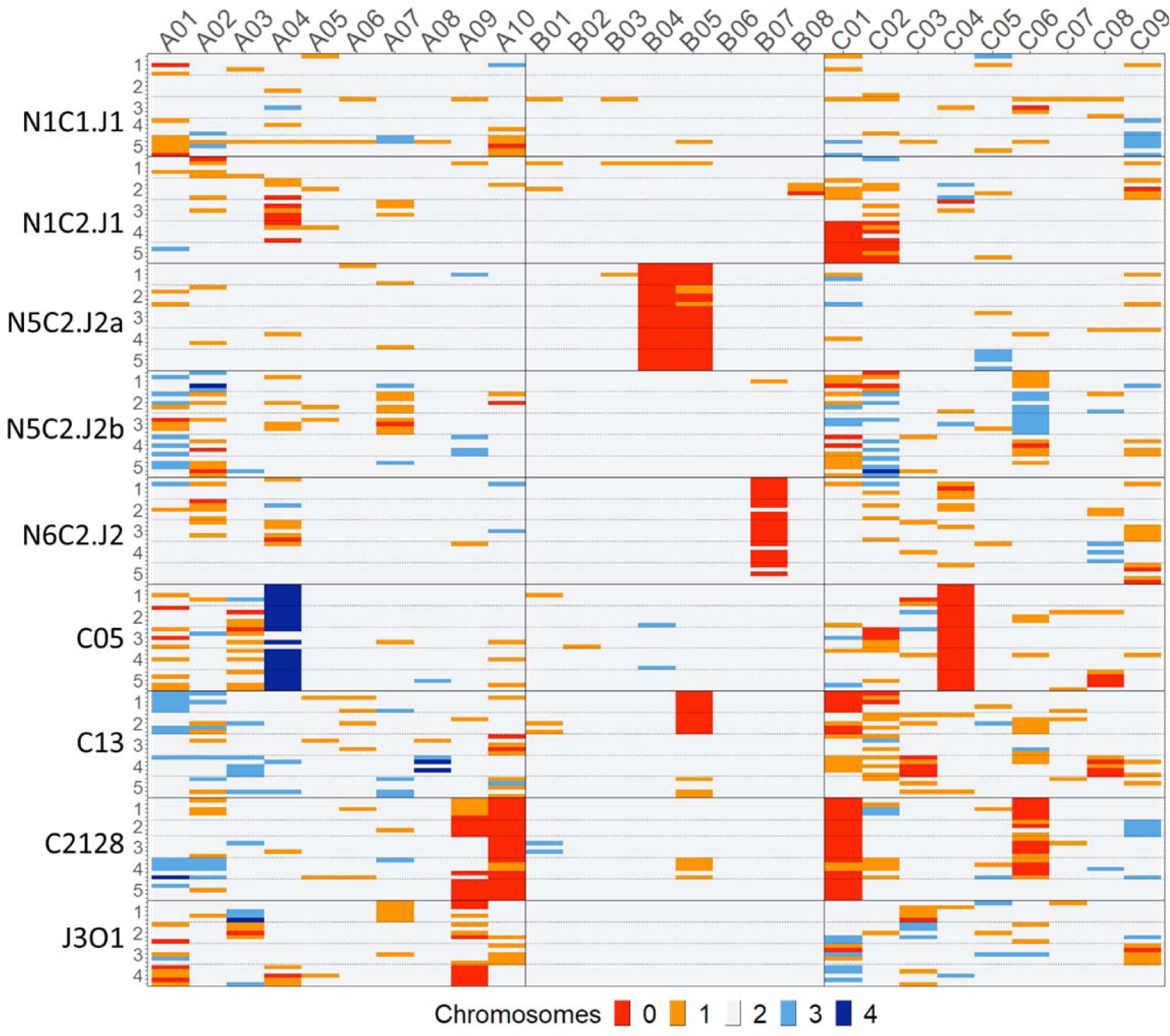
Chromosome copy number variants (CNVs) in different *Brassica* allohexaploid types. Genotype codes are written on the y-axis and within the graph they are marked by a horizontal line. Sibling lines are represented by numbers (1-4 or 1-5) and are marked within the graph by a horizontal dotted line. Within each sibling line 4-5 plants were analyzed, each represented by a line. The number of chromosomes present is colored according to the legend (0 = red, 1 = orange, 2 = light gray, 3 = pale blue and 4 = dark blue) for all chromosomes in subgenomes A (A01 - A10), B (B01 - B08), and C (C01 - C09).

#### Chromosome number in Brassica hexaploids

Nine different genotypes from four different allohexaploids types were selected: NCJ genotypes N1C1.J1, N1C2.J1, N6C2.J2, N5C2.J1a, and N5C2.J1b; carirapa genotypes C05, C13, C2128; and junleracea genotype J3O1. Out of the 218 allohexaploid plants analyzed, 20 had the expected chromosome number of 2n = 54, but only seven were true euploids with the correct numbers of chromosomes in each subgenome. The N1C1.J1 lineage contained the majority of the euploid plants (71%); N1C1.J1 line 2 from this genotype had 3/5 individuals with a euploid chromosome number, although with fixed and segregating chromosome rearrangement events (Figure 3). Different CNVs affecting chromosome number were also identified, with 0 to 4 copies present for each chromosome (Figure 2).

**Figure 3.**
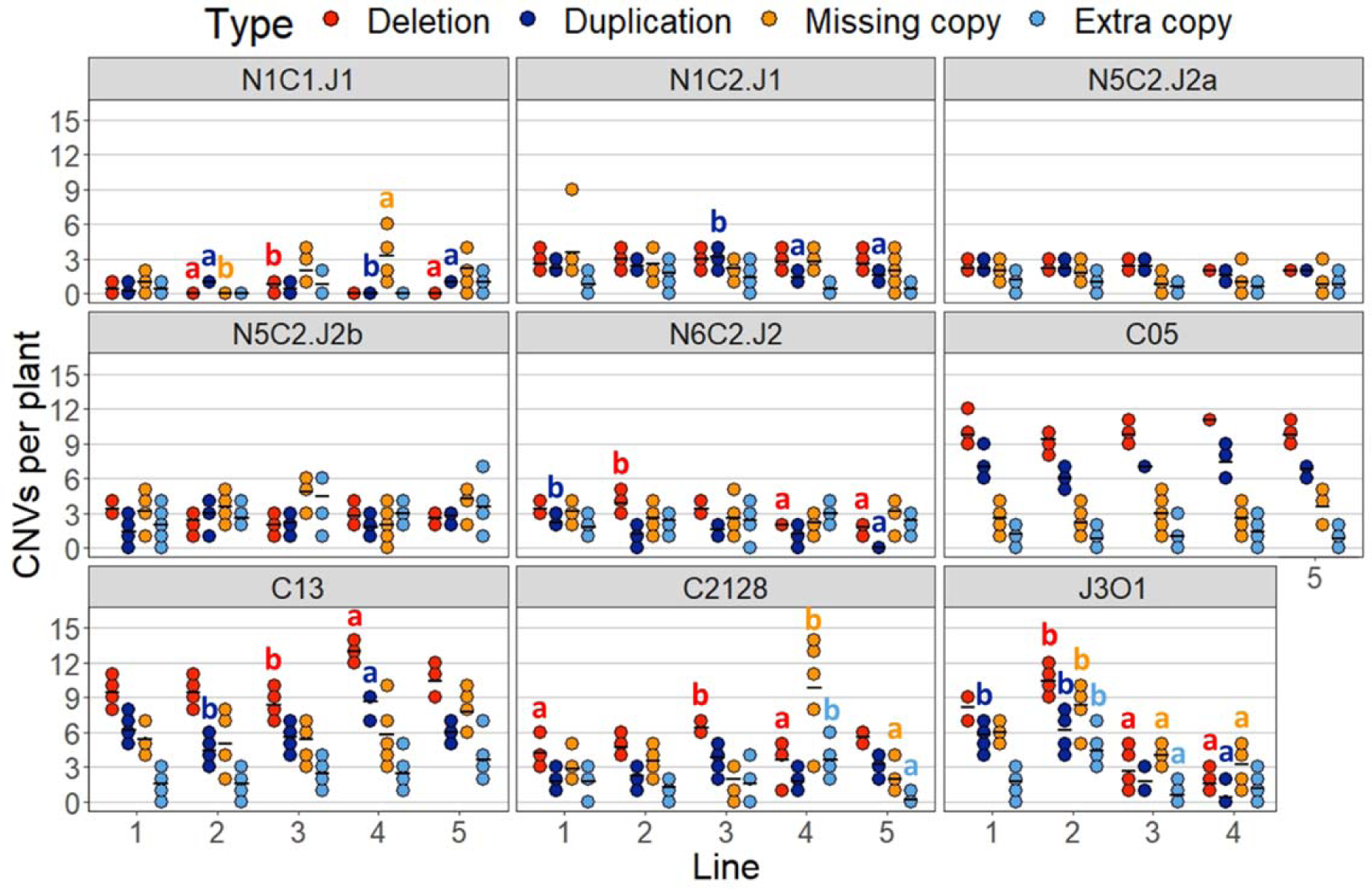
Copy number variants (CNVs) in *Brassica* allohexaploid lines grouped by genotypes. NCJ (*B. napus* × *B. carinata* × *B. juncea*) genotypes: N1C1.J1, N1C2.J1, N5C2.J2a, N5C2.J2b, and N6C2.J2; carirapa (*B. carinata* × *B. rapa*) genotypes: C05, C13, and C2128; junleracea (*B. oleracea* × *B. juncea*) genotype J3O1. Comparisons were made per CNV type between the lines of the same genotype. Each CNV type is colored according to the legend: red indicates deletions (zero copies), dark blue indicates duplications (four copies), orange indicates a missing copy (one copy) and pale blue indicates an extra copy (three copies) of each chromosome region. Mean values per line are shown as a horizontal black line. Statistically significant differences between the lines are shown with letters (the same color of letter represents the same CNV type comparison), where different letters represent significant differences between the lines (Kruskal-Wallis test, followed by Dunńs test, significance at *p* < 0.05).

#### Chromosome number variants in the NCJ allohexaploid type

In the NCJ genotypes, 114 events involving the loss of both chromosome copies, 194 events involving one missing copy, 73 events involving an extra chromosome copy, and two events involving two extra copies of a chromosome were identified. In the NCJ genotypes, most of the events involving zero copies of a chromosome were present in the B genome, as seen in genotype N5C2.J2a for chromosomes B04 (all individuals with 0 copies), B05 (22 individuals with 0 copies and three individuals with one copy) and in genotype N6C2.J2 for chromosome B07 (20 individuals with 0 copies) (Figure 2). Single-copy chromosomes were frequently observed, especially for chromosome A02 (21 events) affecting mostly genotypes N5C2.J2b (seven plants from different lines), N6C2.J2 and N1C2.J1 (six plants each from different lines); and for chromosome C01 (19 events, mostly in genotype N5C2.J2b - 9 plants from different lines). Extra copy chromosomes in NCJ types were also observed particularly for chromosomes A01 and C02 (10 events in total for each) in genotype N5C2.J2b (in eight plants from different lines). Two extra copies of a chromosome were only observed in two instances in the NCJ allohexaploids, with one event each involving chromosomes A02 and C02 respectively, in one plant each of genotype N5C2.J2b lines 1 and 5. For both of these events the corresponding homeologous chromosome had zero copies (centromere was deleted) but the remaining regions of the chromosome were still present and translocated into the two extra copies (Figure 2).

#### Chromosome number variants in the carirapa allohexaploid types

In the carirapa genotypes, there were 131 completely absent chromosomes (zero copies), 137 instances of one chromosome copy, 60 instances of three chromosome copies, and 28 instances of four chromosome copies. Here, unlike the NCJ type, most of the chromosomes with zero copies were located in the C subgenome: in chromosomes C01 (29 observations) with the majority of these in only one genotype C2128 (22 out of 24 individuals analyzed), and in chromosome C04 (25 observations), present in all the plants analyzed from the genotype C05 (Figure 2). Interestingly, the homoeologous chromosome of C04, chromosome A04, was fixed and doubled in 22 of the 25 plants and the remaining region of C04 was translocated into the two extra copies of the chromosome (bottom 12.3 Mb of the chromosome). Single-copy missing chromosomes in carirapa types were observed in the highest number for chromosome C02 (19 observations) present in different individuals from genotypes C05 (four plants), C13 (10 plants), and C2128 (five plants), while an extra chromosome copy was found mostly for chromosome A01 in genotype C13 (eight plants), followed by A02 in genotypes C13 (six plants) and C2128 (four plants).

#### Chromosome number variants in the junleracea allohexaploid type

Junleracea genotype J3O1 had overall fewer chromosome CNV events compared to NCJ and carirapa allohexaploids, with only 16 observations of zero chromosome copies, 45 observations of one chromosome copy, 18 observations of three chromosome copies, and only one observation of two extra chromosome copies (chromosome A03). Most of the zero copy observations were present for chromosome A09 in lines 1 (three plants) and 4 (five plants), while single copy chromosomes were more evenly distributed between different chromosomes. Three copies of a chromosome were mostly observed for chromosome C01 in lines 2 (two plants), 3 (one plant), and 4 (three plants).

#### Genomic stability ranking between allohexaploid types based on segregating chromosome variation

Despite an overall high frequency of karyotype changes, some lines showed little chromosome variation across the five plants analyzed. In order to assess if karyotype change and hence genomic instability was still ongoing, segregating events were counted, since these events are more likely to indicate novel events, and hence may indicate if the karyotype has stabilized or not. Based on this metric, the 44 lines analyzed were ranked in terms of segregating CNV events (Supplementary Figure 1). From 1 up to 28 events per line were identified. Based on this analysis, the most stable genotype was N5C2.J2a line 3, which had only one single missing chromosome event involving C05 in one plant, although chromosomes B04 and B05 were completely lost in this genotype (fixed deletion/loss). The next most stable line based only on chromosome number changes was genotype N1C1.J1 line 2, which as mentioned previously was the line with the most euploid plants. This line also had only two segregating whole chromosome events present in two plants (one missing a copy of A04 and one missing a copy of C02). Carirapa C2128 line 5 also had few changes in single copy chromosomes, with only two plants affected: one with an extra chromosome A01 and one with a missing chromosome A02. This line had also completely lost chromosomes A09, A10, and C01. Interestingly, the line that had the highest number of segregating chromosome changes was also from carirapa C2128 line 4, with 28 events found in four of the five plants analyzed. The junleracea genotype was very unstable compared to many NCJ and carirapa lines, where the most stable line from this genotype was J3O1 line 4 with 14 single chromosome CNVs.

### CNV and karyotype rearrangements in *Brassica* hexaploids

#### *Fixed rearrangement events inherited from the parent allotetraploid species* B. napus, B. carinata, *and* B. juncea

We analyzed the total number of rearrangements in each of the *Brassica* allohexaploid lines per genotype. We identified pre-existing genomic rearrangements in *Brassica* parental genotypes N1, N5, and C1 (Figure 1). Also, from a previous study we identified rearrangements in the parental genotype N6 (deletion on chromosome C02, located between 8-10.7 Mb) (Quezada-Martinez et al. 2022). To avoid bias when analyzing the number of CNVs present we looked at the corresponding allohexaploid produced by crosses of these genotypes, and only 84 fixed events (61 deletion and 23 duplications) were counted as directly inherited from the parents. These rearrangements were mostly present in genotype N1C1.J1, which had 46 events in total in the 24 plants analyzed (22 plants had a fixed deletion of the top of C01 and 24 plants had a fixed deletion of the bottom of C09), and in 15 individuals from genotype N1C2.J1 lines 1, 2, and 3 (14 plants with a fixed deletion of the top of C01 and one plant with a fixed deletion of the bottom of C09). The inherited fixed duplication events were only present in the allohexaploid genotype N1C1.J1, where the bottom of chromosome A10 was doubled and translocated into chromosome C09 in 23 of the 24 plants analyzed. We removed these inherited events from further analysis and kept only events characterized as new.

#### Genomic localization and distribution by allohexaploid type of fixed deletion and duplication events not involving changes in chromosome number

In total, we identified 1002 fixed deletion events and 664 fixed duplication events distributed between the allohexaploid types present on different chromosomes, where these events did not affect chromosome number. These events did not affect chromosome number because they were located in distal chromosome regions, i.e. not spanning or involving the centromere, and we used presence of the centromere as a proxy for chromosome number. The chromosome with the highest number of fixed deletions was C01, accounting for 12.9% of the total, followed by chromosome A04 (9.2%), and chromosome A07 (9%). Six chromosomes (B01, B03, B04, B05, B06, and B08) did not show any fixed deletion events. The chromosome with the most fixed duplication events was C04, accounting for 13.5% of the total, followed by chromosomes C06 (10.8%), C02, and A01 (10.5%). Most of the chromosomes that did not have fixed duplication events were also from the B subgenome (B02, B03, B04, B07, B08), and one chromosome was from the C subgenome (C07). Most of the fixed deletion events were in a size range below or equal to 10 Mb (86.8%, Supplementary Figure 2) with an overall average size of 5 Mb and a minimum size of 1 Mb, the minimum size we were able to assess using this analysis method, and a maximum size of 26 Mb, equivalent to the loss of the bottom part of chromosome C07 in three individuals of carirapa allohexaploid genotype C13 line 5. Similarly, most of the fixed duplication events were ≤ 10 Mb (88.7%, Supplementary Figure 2) with an average of 5 Mb, a minimum size of 1 Mb and a maximum size of 35 Mb (a duplication at the top of chromosome C02 in one plant of N6C2.J2 line 3).

The size of fixed deletions and duplications differed significantly between allohexaploid types (Kruskal-Wallis test, *p =* 1.28e-19), with NCJ deletions being significantly larger (average of 7 Mb) than carirapa (Dunńs test, *p =* 1.07e-19, average of 4 Mb) and junleracea (Dunńs test, *p =* 1.38e-2, average of 5 Mb), and carirapa deletions being significantly smaller than junleracea type (Dunńs test, *p =* 1.45e-3, Supplementary Figure 3). Fixed duplication size also differed significantly between allohexaploid types (Kruskal-Wallis test, *p =* 1.42e-6) with NCJ types showing significantly larger deletions than carirapa types (Dunńs test, *p =* 6.85e-7, Supplementary Figure 3).

#### Genomic distribution of segregating CNV events in the allohexaploids

The number of segregating translocations was quantified in the *Brassica* allohexaploids. Eight events were removed based on parental inheritance: five missing copies of chromosomes A01, B01, and A04, and three duplications in C04: these were present in six plants of the genotype N1C1.J1. In total, 710 events involving the loss of a copy and 359 events involving the gain of a copy in the terminal regions of chromosomes were identified. Missing single copy events were evenly distributed between the A and C subgenomes, with 48.3% and 48.4% of the events, respectively, while the B genome had only 3.3% of the events. In the case of extra copies, the A subgenome (45.5% of events) and C subgenome (60.7% of events) were not significantly different (Pearson’s χ^2^ test, *p =* 0.137), and the B subgenome had 1.4% of these events. C01 had the most missing regions, with 107 events (15.3%), followed by A01 with 70 events (10%), and in third position chromosome C02 with 61 events (8.7%). Chromosome C02 had the most duplication events, with 62 (17.3%), followed by chromosome A09 with 36 events (10%), and chromosome A01 with 35 events (9.7%).

#### Fixed deletion and duplication events not involving changes in chromosome number varied between lines within genotypes of NCJ, carirapa, and junleracea allohexaploid types

In NCJ allohexaploids, 268 fixed deletion events, 206 fixed duplication events, 288 missing copy events, and 194 extra copy events were identified. The different lines per genotype and the number of events per plant were compared to assess the effect of independent rearrangements segregating in the genotypes. Lines within the genotypes N5C2.J2a and N5C2.J2b had similar number of events per plant with no significant differences in any of the events analyzed (fixed deletions per plant, Kruskal-Wallis test, p *=* 0.406 and *p =* 0.0846; fixed duplications, Kruskal-Wallis test, *p =* 0.118 and *p =* 0.0894; missing copy, Kruskal-Wallis test, *p* = 0.157 and *p* = 0.0536; and extra copy, Kruskal-Wallis test, *p* = 0.623 and *p* = 0.383, respectively; Figure 3). Lines in the genotype N1C2.J1 only differed significantly in the number of duplication events per plant (Kruskal-Wallis test, *p* = 0.00757). Lines in the genotype N6C2.J2 only differed significantly in the number of fixed events per plant: deletions (Kruskal-Wallis test, *p* = 6.5e-4) and duplications (Kruskal-Wallis test, *p* = 5.4e-3). In the genotype N1C1.J1 the sibling lines had more variation and significant differences were observed in three categories: deletions (Kruskal-Wallis test, *p* = 0.0124), duplications (Kruskal-Wallis test, *p* = 0.00415), and missing copy (Kruskal-Wallis test, *p* = 0.0144).

In carirapa allohexaploids, 620 fixed deletions and 387 fixed duplication events were identified. Lines in the carirapa genotype C05 showed no significant differences in any of the rearrangement types analyzed (deletions, Kruskal-Wallis test, *p =* 0.0758; duplications, Kruskal-Wallis test, *p* = 0.188; missing copy, Kruskal-Wallis test, *p =* 0.594; and extra copy, Kruskal-Wallis test, *p* = 0.826; Figure 3). Lines in the genotype C13 showed only significant differences in the fixed events: deletions (Kruskal-Wallis test, *p* = 4.7e-3) and duplications (Kruskal-Wallis test, *p* = 6.5e-3). On the other hand, lines in the genotype C2128 varied significantly, with statistical differences found in all the rearrangement types analyzed: deletions (Kruskal-Wallis test, *p* = 5.8e-3), duplications (Kruskal-Wallis test, *p* = 0.0365), missing copy (Kruskal-Wallis test, *p* = 0.0243), and extra copy (Kruskal-Wallis test, *p* = 0.0164) events.

Finally, in the junleracea allohexaploids (single genotype J3O1) 114 fixed deletion events and 71 fixed duplication events were found. Significant differences between the lines were found for all CNV types: deletions (Kruskal-Wallis test, *p =* 0.00101), duplications (Kruskal-Wallis test, *p =* 0.0013), missing copy (Kruskal-Wallis test, *p* = 3.1e-3), and extra copy (Kruskal-Wallis test, *p* = 0.0112, Figure 3) events.

#### Duplication events without an identifiable genomic location for the duplicated region

For some single copy duplication events the specific location in the genome could not be determined, either because there was no missing copy in the respective homeologous region or because no missing copy in the rest of the genome was scored according to the parameters (> 1 Mb in size). In the NCJ allohexaploids, 26 duplications of A-genome chromosome regions (A01, A07, A09, and A10) with unknown chromosomal location were identified. Most of these events (19) involved a duplication of the bottom part of A10 (located between 18.4 – 19.9 Mb) in genotype N6C2.J2 in different lines (line 1 - four plants, line 2 - five plants, line 4 - five plants, and line 5 - five plants). In the carirapa genotypes, only two single copy duplication events were identified of chromosome A09 regions with unknown locations, one located at the top of the chromosome (0 - 3 Mb) and one at the bottom of the chromosome (43 – 46.7 Mb), present in the same genotype of C2128 in line 4, but in two different plants. In the junleracea allohexaploids a single duplication event was found with an unknown location in chromosome A10, located between 0 - 1.9 Mb in a plant of line 2. Only one single copy duplication event with an unknown genomic location was observed in the B genome: this was in a carirapa plant from genotype C13 line 4, where B05 had a single copy duplication of the region from 13.8 – 23.4 Mb. For the C genome, 30 single copy duplication events were identified in NCJ allohexaploids, with most of them located on chromosome C02 (20 events), while the remaining duplicated regions involved chromosomes C04 and C09 (four events each), and C07 and C08 (one event each). In the carirapa type eight events were found, with three events located on chromosomes C01 and C03, and one single event for each of chromosomes C02 and C07. For the junleracea five duplication events were identified, with unknown genomic location: two events located on chromosome C05, and one event in each of chromosomes C01, C02, and C03.

#### Differences between genotypes in frequencies of different types of CNVs and correlations with selfing-rounds

Overall, significant differences were observed between genotypes for all CNV types analyzed: deletions (Kruskal-Wallis test, *p =* 1.29e-31), duplications (Kruskal-Wallis test, *p =* 1.71e-26), missing copy (Kruskal-Wallis test, *p =* 1.58e-15), and extra copy (Kruskal-Wallis test, *p =* 3.67e-12) (Figure 4). Genotype N1C1.J1 had significantly less deletions compared to the other genotypes (Dunńs test, *p* < 0.05, Figure 4).

**Figure 4.**
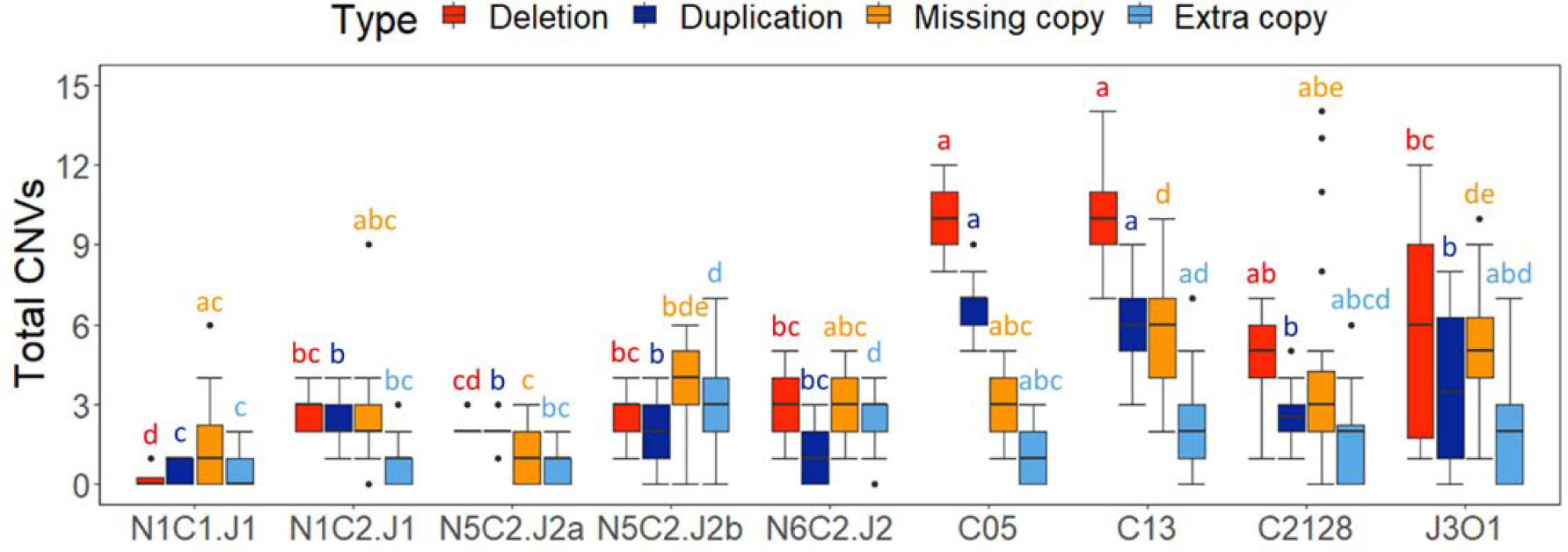
Copy number variants (CNVs) per genotype in different *Brassica* allohexaploids. NCJ genotypes: N1C1.J1, N1C2.J1, N5C2.J2a, N5C2.J2b, and N6C2.J2; carirapa genotypes: C05, C13, and C2128; junleracea genotype J3O1. Comparisons were made per type of CNV between all the allohexaploid genotypes. Each CNV type is colored according to the legend: red indicates deletions (zero copies), dark blue indicates duplications (four copies), orange indicates a missing copy (one copy) and pale blue indicates an extra copy (three copies) of each chromosome region. Statistical differences between the genotypes are shown with letters (the same color of letter represents the same CNV type comparison), where different letters represent significant differences between the genotypes (Kruskal-Wallis test, followed by Dunńs test, significance at *p* < 0.05).

Since we have different generations, we were interested if there was any correlation between the total number of CNVs and the allohexaploid genotype generation: we identified a weak correlation between the two values (*r* = 0.3, *p =* 2.9e-09), where the older generations have more CNVs, as expected. On the other hand, when analyzed separately per allohexaploid type, the NCJ allohexaploids showed a significant negative correlation (*r* = -0.17, *p =* 0.021), and the carirapa allohexaploids showed a significant positive correlation in number of CNVs relative to number of generations (*r* = 0.28, *p =* 0.0019).

### Effects of genotype and allohexaploid type on total CNVs

Allohexaploid type on its own (NCJ, carirapa, or junleracea) was able to explain 50% of the variation observed in total CNV numbers: carirapa, junleracea and NCJ allohexaploid type had effects of +24.3, +20.7, and +10.9 CNVs per plant (*p* = <2e-16, *p* = 0.0325, and *p* = <2e-16, respectively). Genotype-specific effects were also calculated in a linear model that was able to explain 70% of the variation observed in CNV accumulation, with all genotypes analyzed having a significant effect on total CNV accumulation (Supplementary Table 5). Significant effects of genotype on CNV totals ranged from +5.5 to +29.4 CNVs per plant. The smallest positive effects were found in genotypes N1C1.J1 (+5.5 CNVs, *p* = 4.3e-14), N5C2.J2a (+9.3 CNVs, *p* = 1.82e-4), and N1C2.J1 (+11.5 CNVs, *p* = 3.85e-08), while the largest positive effect was observed for genotypes C13 (+24.9 CNVs, *p* = <2e-16), C05 (+24.9 CNVs, *p* = <2e-16), and J3O1 (+20.7 CNVs, *p* = <2e-16).

NCJ allohexaploids were a combination of a few genotypes from *B. napus* (N1, N5, and N6)*, B. carinata* (C1 and C2), and *B. juncea* (J1 and J2). In the NCJ set we analyzed for the independent effect of these parental genotypes on the total number of CNVs. *Brassica napus* genotypes showed a positive effect on the total CNVs but only for two genotypes, with genotype N1 having the lowest effect of +8.5 total CNVs per plant (*p* = 1.21e-05), followed by N5 with +12.5 total CNVs per plant (*p* = <2e-16). *Brassica carinata* genotypes had more significant contrasting effects, with C1 having an effect of +5.5 (*p* = 1.48e-10), while C2 had an effect of +12.2 (*p* = 3.78e-12) total CNVs. Finally, *B. juncea* genotypes had a significant effect on total number of CNVs per plant of +8.5 (*p* = <2e-16) and +12.4 (*p* = 7.38e-06) for genotypes J1 and J2, respectively. Overall *B. napus, B. carinata,* and *B. juncea* on their own explained less than 35% of the variation observed (14%, 32%, and 15%, respectively).

### Translocation events and subgenome bias in *Brassica* allohexaploids: fixed duplication/deletion events

For all fixed duplications not involving centromeric regions and hence most likely to comprise non-reciprocal translocation events, we were able to identify where in the genome the extra copies were located based on inspection of the primary homoeologous region. In total, we found that putative translocation events involving a C fragment replacing an A fragment (256 events) were significantly more common than an A fragment replacing a C fragment (372 events, respectively. Pearson’s χ^2^ test, p = 3.68e-06). Specifically, 12 different chromosome pairs were identified with fixed putative non-reciprocal translocations involving a duplication of the A genome translocated into the C genome (256 events of an A fragment into C genome chromosome). These events involved chromosomes A01 – C01 (10.5% of the total fixed non-reciprocal translocation events), A07 – C06 (9.6%), A09 – C08 (5.7%), A09 – C09 (4.5%), A03 – C03 (4.2%), A02 – C02 (2.4%), A10 – C09 (0.6%), A04 – C04 (0.3%), A05 – C04 (0.2%), A05 – C05 (0.2%), A06 – C05 (0.2%), and A07 – C07 (0.2%). Twelve different chromosome pairs were also involved in a fixed translocation of a C genome fragment into an A genome chromosome (372 events): C04 – A04 (11.7% of the total fixed non-reciprocal translocation events), C06 – A07 (10.8%), C02 – A02 (10.5%), C03 – A03 (6.2%), C09 – A09 (4.2%), C05 – A05 (3.8%), C01 – A01 (2.6%), C04 – A05 (1.8%), C08 – A09 (1.7%), C05 – A06 (1.5%), C09 – A10 (1.1%) and C08 – A08 (0.2%).

The greatest number of putative translocations was observed between C04 and A04, where the C region was doubled and translocated into the A subgenome: these accounted for 78 events in total, with all these events involving the bottom portions of the chromosomes. At the same time, when the bottom of C04 was deleted (11 events), in only two events was the corresponding homoeologous region in A04 doubled and putatively translocated. For the B genome, only identified four different fixed putative non-reciprocal translocations were identified, all involving translocation of a B genome fragment into a C genome chromosome): B01 – C05 (3.6% of the total non-reciprocal translocation events), B05 – C01 (0.8%), B01 – C04 (0.6%), and B06 – C06 (0.5%). The size of the B genome fragment translocated varied, from 1.4 Mb (B05→C01 in two plants in C13 line 5) up to 17.6 Mb (B01→C04 in four plants of J3O1 line 1).

In the NCJ type, the number of A → C and C → A subgenome translocations were approximately the same (∼50%), with no translocations involving B chromosomes. In carirapa allohexaploids, there were significantly more translocation events (59.9%, Pearson’s χ^2^ test, *p =* 2.12e-8) that involved a C chromosome segment replacing an A chromosome segment, while only 32.6% of events involved translocation of an A chromosome segment into the C genome, and 7.5% of events involved translocations from the B into the C genome. In the junleracea hexaploids, 47.9% of events involved translocation from the C to A genome, 42.3% involved translocation from the A to C genome, and 9.9% involved translocation from the B to C subgenome. Genotypes and lines within genotypes also varied in direction of fixed translocations between subgenomes (Figure 5).

**Figure 5.**
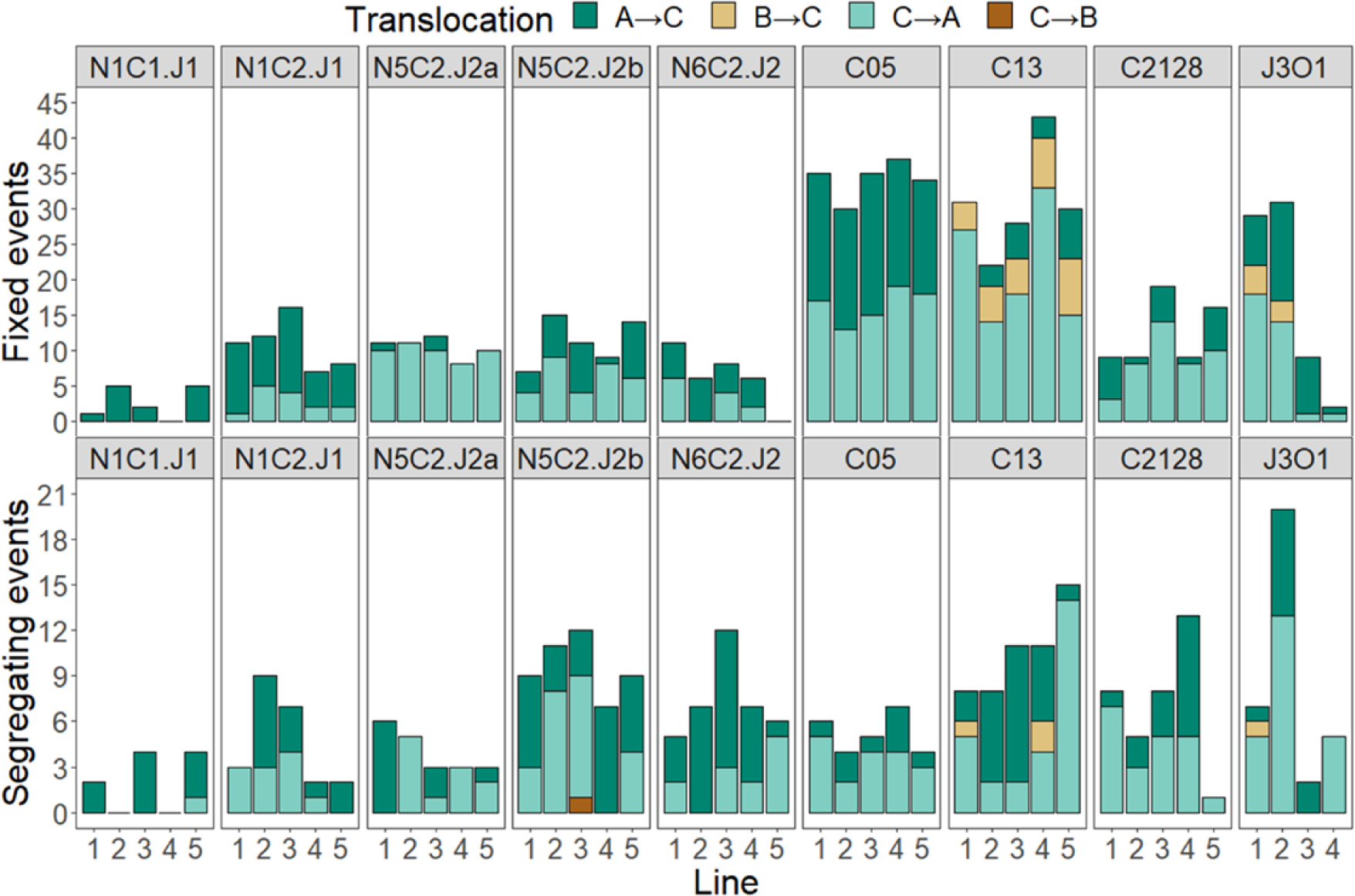
Translocations between the different *Brassica* subgenomes (A, B and C) in different allohexaploid genotypes. NCJ type (*B. napus* × *B. carinata* × *B. juncea*) genotypes: N1C1.J1, N1C2.J1, N5C2.J2a, N5C2.J2b, and N6C2.J2. Carirapa type (*B. carinata* × *B. rapa*) genotypes: C05, C13, and C2128. Junleracea type (*B. juncea* × *B. oleracea*) genotype J3O1. Genotype names are at the top of each rectangle. The upper graph represents the number of fixed translocations between the different subgenomes (see legend: e.g. A → C = A-genome chromosome fragment duplicated and translocated into the C subgenome). Each bar represents one line per genotype (5 lines for the NCJ and carirapa allohexaploid types and 4 lines for the junleracea type).

### Translocation events and subgenome bias in *Brassica* allohexaploids: segregating events (single copy / extra copy translocations)

A total of 286 single-copy (heterozygous, segregating) homoeologous translocation events between subgenomes were identified (Figure 5). The chromosomes commonly involved in non-reciprocal translocation events was similar between allohexaploid types, although the direction of the event differed, with no significant bias overall towards the direction of the translocation between the A and C subgenomes (Pearson’s χ^2^ test, *p* = 0.438). In the NCJ allohexaploid type, 138 events involving a putative translocation were found, where most of the events corresponded to a duplicated A chromosome fragment translocated into a C-genome chromosome (57.2%), or to a duplicated C-genome chromosome fragment translocated into an A-genome chromosome (42%). Only one putative translocation involved the B genome, where a fragment of chromosome C04 was translocated into B06. In the carirapa allohexaploid type, most of the translocations occurred in the opposite direction from the NCJ, and involved a duplication of a C-genome chromosome fragment translocated into an A-genome chromosome (57.9%), while the A → C direction accounted for 39.5% of the events, and B chromosome fragments translocated into C-genome chromosome comprised 2.6% of the events. Finally, in the junleracea type, as in the carirapa type, most events involved translocation from the C subgenome into the A subgenome (67.6%): A → C events only accounted for 29.4% of the total, and one instance of a B-genome chromosome translocation into the C subgenome was observed between chromosomes B01 and C04.

### Crossing *Brassica* allohexaploid genotypes

Flowering time varied between genotypes (Supplementary Table 2): the shortest flowering time was 32 DAS (one plant in genotype N4C2.J1), while the longest was 121 DAS (two plants in genotype N5C2.J2a), which affected the order in which plants were crossed. Genotype N5C2.J2a had the widest range of flowering time with an average of 95 ± 23 DAS, while the earliest flowering genotype N1C2.J1 had the least variation (average of 43 ± 1 DAS). Pollen viability for each plant was also estimated: 62% of the plants had < 50% viable pollen, and only 6% had pollen viability > 80% (Supplementary Figure 4). Two naponigra genotypes (N8.I1 and N9.I2) had poor anther development and produced no viable pollen (0%). Similarly, in the case of the N5.I2 naponigra genotype, pollen viability estimates could only be obtained from two of the four plants available.

In total, 113 different cross-combinations were carried out with 10 310 flower buds crossed. Out of the total crosses harvested, 48.7% (5023 crosses) developed into siliques, with a maximum success rate of 97.1% silique development in a cross between N7C1.J1 and N1C2.J1, where only 34 flower buds were crossed. However, most of the crosses produced a low ratio of seeds per bud pollination, ranging from 0.0 – 4.6 seeds/flower bud (Figure 6).

**Figure 6.**
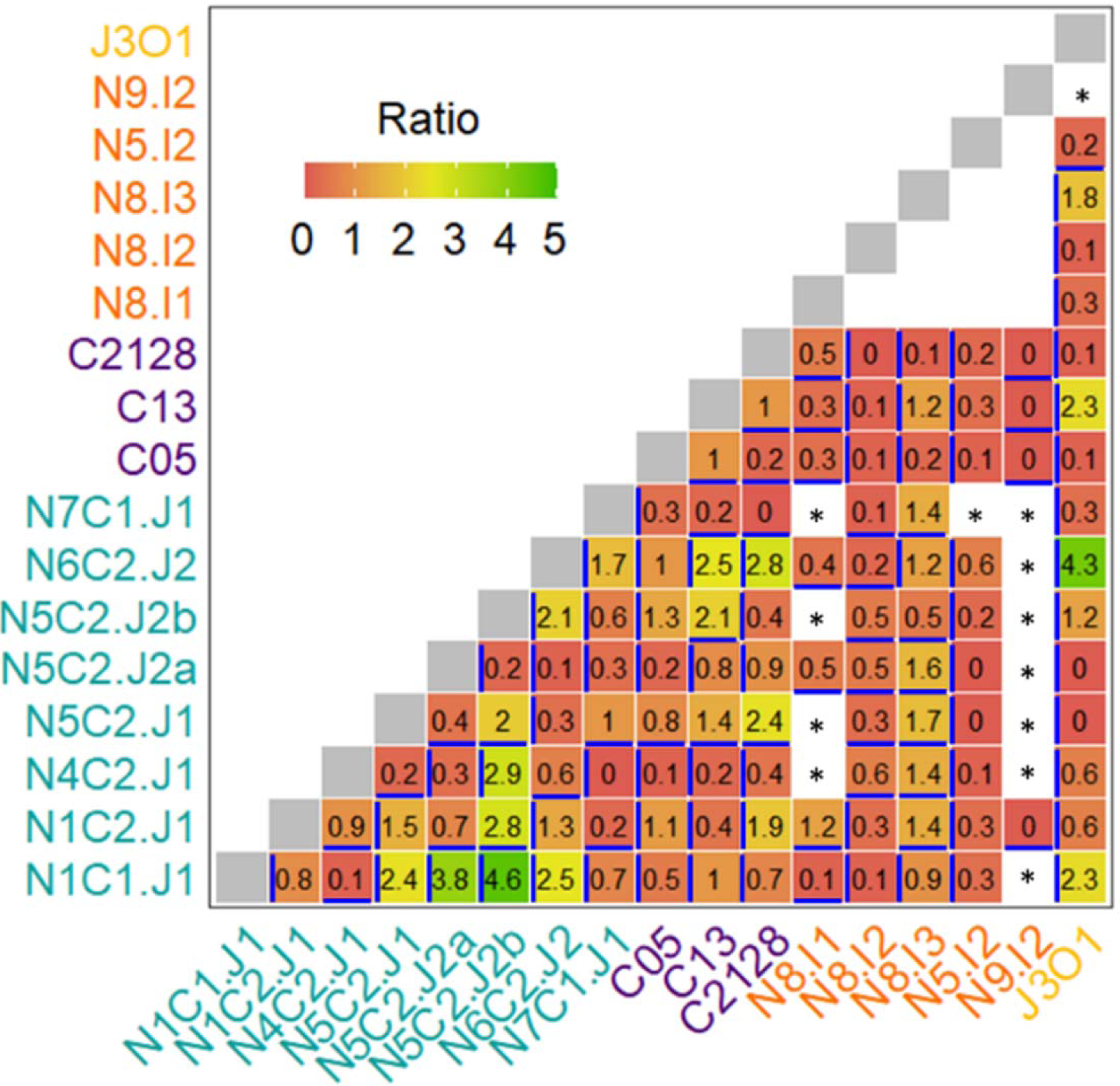
Crossing success between different allohexaploid *Brassica* genotypes. Ratio represents the total number of seeds produced divided by the number of flower buds cross-pollinated between genotypes. * Cross combination was planned but not performed due to experimental constraints. The female parent in the cross is marked with a perpendicular blue line relative to the genotype name: e.g. in N1C1.J1 × N1C2.J1, the female parent is N1C1.J1.

For some of the crosses, siliques were developed, but upon opening contained only shriveled seeds or no seeds at all. This was observed especially when N5.I2 was used as a male donor in the cross. We identified at least four cross combinations where 50% of the siliques developed after crossing, but very few seeds were obtained (a ratio lower than 0.3 seeds/bud pollination): genotype combinations N5C2.J2a × N5.I2, J3O1 × N5.I2, C2128 × N5.I2, and N5C2.J2b × N5.I2. The maximum number of seeds produced from a cross was obtained from N1C1.J1 × N5C2.J2a, where out of 109 flower buds crossed, 500 seeds were obtained (Ratio = 4.6, Figure 6).

Genotype affected the ratio of seeds produced per bud pollination (Supplementary Table 6). Genotype N1C1.J1 had a strong positive effect on the success ratio of seeds/bud pollination when used as female parent, adding an extra 1.0 seeds per bud pollination based on multiple regression analysis (*p =* 0.00431). Seven other genotypes showed a positive effect on ratio of seeds produced per bud pollination, ranging from 0.01 seeds/bud in genotypes N7C1.J1, up to 0.28 seeds/bud in genotype C05 (Supplementary Table 6). Genotype N9.I2 showed a negative effect as a female parent, with -0.32 effect in seeds/buds. Male parent genotype effect was only significant for one genotype (two lineages) with big effect of +1.82 and +1.95 observed for N5C2.J2a and N5C2.J2b. Together, female and male genotype combination was able to explain 47% of the variation observed in the ratio of seeds produced per cross combination (Supplementary Table 6, Figure 6).

Crossings between NCJ genotypes provided an excellent opportunity to analyze the effect of having a common *Brassica* parent genotype on both sides of the cross on the number of seeds produced per bud pollination (ratio, Supplementary Table 7). For example, in the cross combination N4C2.J1 × N5C2.J2a, *B. carinata* genotype “94024.2_02DH” (C2) is in common between the maternal and paternal parent line. Based on multiple linear regression analysis, having one (*p =* 0.1642) or two parents (*p =* 0.0952) in common had no significant effect on the ratio of seeds produced per bud pollination. Interestingly, having zero parents in common had a greater effect of +2.0 on seeds/bud pollination (*p =* 0.0470). Having three parents in common had a negative effect of -1.1, but only one cross combination represented this category (*p =* 0.0108). In the same model but considering only NCJ genotypes and including the female and male genotypes separately, only one genotype had a significant effect: genotype N5C2.J2a used as a female parent had a slight positive effect on ratio of seeds produced per bud pollination (+0.6; *p =* 0.0172), and N5C2.J2b used as a male parent had a higher positive effect on ratio of seeds produced per bud pollination (+4.7, *p =* 0.01388).

In total, 9052 new hybrid seeds were obtained from cross-pollinations between allohexaploids, out of which 835 had germinated in the silique at the time of harvesting. On average, 14.2% of the seeds were viviparous per cross combination. Maternal and paternal genotypes were compared in the number of viviparous seeds produced and significant differences were found only when comparing maternal genotypes used in the cross (Kruskal-Wallis test; *p =* 5.7e-07) and not when comparing male parent genotypes (Kruskal-Wallis test; *p =* 0.3713). Using a linear regression model, four genotypes had a significant positive effect on the percentage of viviparous seeds produced: N7C1.J1 (+34.1%), N5C2.J2 (+31.5%), N4C2.J1 (+29.7%), and N1C1.J1 (+13%); however, the maternal genotype only explained 25% of variability observed (Supplementary Table 8).

The overall ratio of self-pollinated seeds produced per bud pollination was higher than the ratio obtained by crossing, ranging from 0.0 – 11.4 seeds/flower bud. Out of the 95 plants used for crossing, 17 were completely self-infertile, and genotype N9.I2 was completely sterile. Out of all the naponigra genotypes, seeds were only obtained from two plants of genotype N8.I2, where five harvested branches (secondary meristems) produced a total of eight seeds. Two of the five *Brassica* junleracea J3O1 genotypes were also self-infertile, together with one plant each of genotypes N7C1.J1 and N4C2.J1.

Total seed number and pollen viability were weakly correlated (*r* = 0.19, *p =* 0.0095), as were ratio of self-pollinated seeds per bud pollination and pollen viability (*r* = 0.25, *p =* 0.00069) and ratio of self-pollinated seeds per bud pollination and flowering time (*r* = 0.25*, p =* 0.0017). A stronger correlation was observed between number of developed siliques and total number of seeds obtained (*r* = 0.7, *p =* <2.2-16).

### F_1_ hybrids compared to the parents

Eight F_1_ hybrids were selected based on the number of seeds obtained and the different parental genotype combinations involved (Table 1). Unfortunately, self-pollinated seeds from one of the parents of the hybrids could not be obtained, genotype N8.I3, and hence this parent line is missing as a control to compare with the new F_1_ hybrids. All the F_1_ hybrids analyzed were true hybrids between the parents based on the genotyping data analysis.

**Table 1.** Crosses between *Brassica* hexaploids NCJ (*B. napus* × *B. carinata* × *B. juncea*), junleracea (*B. juncea* × *B. oleracea*), carirapa (*B. carinata* × *B. rapa*), and naponigra (*B. napus* × *B. nigra*) genotypes selected for comparison with the parents. The individual parental plant used in the cross is represented by a hyphen followed by the number of the plant.

| Genotype<br>Female | Genotype<br>Male | Hybrid Genotype | Flower<br>buds<br>crossed | Siliques<br>developed | Seeds<br>obtained | Ratio* |
| --- | --- | --- | --- | --- | --- | --- |
| N1C1.J1-1 | N6C2.J2-2 | N1C1J1 × N6C2J2 | 99 | 58 | 251 | 2.5 |
| N6C2.J2-1 | C13-1 | N6C2J2 × C13 | 110 | 62 | 277 | 2.5 |
| N6C2.J2-1 | J3O1-1 | N6C2J2 × J3O1 | 44 | 41 | 191 | 4.3 |
| N6C2.J2-3 | N5.I2-1 | N6C2J2 × N5I2 | 66 | 37 | 42 | 0.6 |
| C13-2 | J3O1-1 | C13 × J3O1 | 100 | 79 | 231 | 2.3 |
| C13-2 | C05-1 | C13 × C05 | 100 | 44 | 103 | 1.0 |
| C13-3 | N8.I3-1 | C13 × N8I3 | 87 | 66 | 101 | 1.2 |
| N8.I3-1 | J3O1-2 | N8I3 × J3O1 | 106 | 97 | 186 | 1.7 |

\*Seeds obtained per cross-pollinated flower bud

Out of the eight F_1_ hybrids, only three showed significant differences in flowering time compared to either of the parents (Supplementary Table 9). Hybrid C13 × J3O1 averaged 52 days to flower (DTF), significantly fewer than parent genotype C13-2 (average 78 DAS; Student’s pairwise t-test, *p =* 0.0388), but not parent genotype J3O1-1 (average 45 DAS; Student’s pairwise T-test, *p =* 1), although the parents did differ significantly in flowering time (Student’s pairwise t-test, *p =* 0.0147). Similarly, the new F_1_ hybrid N6C2.J2 × J3O1 flowered significantly earlier (on average 51 DTF) than its parent N6C2.J2-1 (on average 69 DTF, Student’s pairwise t-test, *p =* 0.0392), but no significant difference was observed when compared to the junleracea parent J3O1-1 (Student’s pairwise t-test, *p =* 0.868); flowering time in DAS again differed significantly between the parents (Student’s pairwise t-test, *p =* 0.0105). On the other hand, hybrid N8.I3 × J3O1 took significantly more DAS (72) to flower compared to its parent J3O1-2 (53 DAS; Student’s t-test, *p =* 0.0323). All the other new hybrids showed similar DAS compared to their respective parents.

Four parental plants did not produce any seed in this new generation: J3O1-1, J3O1-2, C05-1, and N5.I2-1. The remaining plants produced between 1 – 3533 seeds (Supplementary Figure 5). Number of seeds produced by F_1_ hybrids compared to their parents differed significantly in three of the cross combination analyzed (Figure 6). Only the F_1_ hybrid C13 × C05 produced significantly more seeds than one of its parents (female parent genotype C13, Figure 6).

### CNVs and seed set in parents and F_1_ progeny

F_1_ hybrids compared to both parents showed no correlation between total CNV present per plant and total number of self-pollinated seeds obtained (*r* = -0.046, *p =* 0.63). However, a correlation was observed between presence of an extra chromosome or extra fragment from the A subgenome and total seed number (*r* = 0.24, *p =* 0.015). Similarly, the number of A-genome chromosomes present and the number of total self-pollinated seeds was also significantly correlated (*r* = 0.21, *p =* 0.031). All the plants analyzed in the F_1_ were aneuploid (Supplementary Table 8), with chromosome number ranging from 43-57 chromosomes in total. Only one F_1_ hybrid differed significantly from either of its parents in terms of number of CNVs present: N8I3.J3O1, where the F_1_ hybrid had fewer CNVs compared to J3O1-2 (Supplementary Figure 6).

One dwarf sterile plant was observed in the parental genotype J3O1-1. During germination, this plant emerged with four cotyledons, and despite being able to develop and flower it did not phenotypically resemble the other sibling lines. This plant did not have a high number of CNVs (16 CNVs) and the majority of the rearrangements affected the number of chromosomes, and as a consequence only 43 chromosomes were present (Supplementary Table 8). Another sterile dwarf plant from the same allohexaploid combination was an individual from J3O1-2, with many rearrangements (39 CNVs) and 47 chromosomes present.

Interestingly, in genotype C05, which only had two plants available, a similar number of CNVs was observed between the plants (27 and 31), but the plant with a lower number of CNVs produced 2223 seeds compared to zero seeds produced by the hybrid with 31 CNV events. CNV events between these plants not only affected chromosome segments, but also chromosome number, where the hybrid that produced more seeds had 53 chromosomes (missing both copies of C04 and one copy of A10, but had 4 copies of A04), and the hybrid which produced zero seeds had 49 chromosomes (missing both copies of A01, A02, and C04, single copy of A07, A10, C07, C08, an extra copy of C01, C02, and C05, and two extra copies of A04). One naponigra plant N5.I2 also produced no seeds, had few CNVs (8) and 52 chromosomes with missing copies of A02, A09, C01 and C06, and an extra copy of C03 and C09. This plant also had no anther development and short filaments.

## Discussion

In this study, we firstly aimed to systematically compare frequencies and types of CNV events observed between advanced allohexaploid *Brassica* lineages produced from different species and genotype combinations. Previous studies have suggested that differences exist between allohexaploid lineages (different species combinations) in terms of meiotic stability (Mwathi et al. 2017) and genomic stability (Zhou et al. 2016), and genotype-specific effects have previously been established to arise from different parents in segregating mapping populations (Gaebelein et al. 2019b), but here we undertake the first comparison across multiple allohexaploid types and genotype combinations, using high-resolution molecular karyotyping. We found strong effects of genotype and allohexaploid type on overall accumulation of CNVs per plant, with carirapa genotypes accumulating the most CNVs, followed by junleracea, then NCJ types. Each of *B. napus, B. juncea* and *B. carinata* parent genotype had a significant influence on total number of CNVs per plant, suggesting genetic factors inherited from each of these species affects non-homologous recombination frequency in the allohexaploids, as also suggested by (Gaebelein et al. 2019b). Similarly, genotypic differences on chromosome pairing behavior (bivalent and univalent frequency during meiosis) have been observed in *Brassica* carirapa allohexaploids, with one parental *B. rapa* genotype identified to confer almost 100% bivalent formation (Gupta et al. 2016). *Brassica* allohexaploids of four different species combinations were previously observed to show some cytological differences in pairing behavior and genomic stability with carirapa (S_7_) having the highest, followed by junleracea S_0-1_, *B. rapa* × *B. oleracea* × *B. nigra* with 44.4% (Zhou et al. 2016). Genotypic effects on the frequency of non-homologous recombination events have also been observed cytologically in *B. napus* (Sheidai et al. 2003), in interspecific hybrids between *B. juncea* and *B. napus* (genome composition AABC; (Mason et al. 2010)), and using molecular marker segregation methods in interspecific hybrids between *B. napus* and *B. carinata* (Mason et al. 2011). In synthetic *B. napus* genotypes, parental *B. oleracea* and *B. rapa* genotype combinations also affected number of CNVs present in progeny after one generation of self-pollination (two meioses) (Katche et al. 2023). Our results suggest that many genetic variants inherited from parent species and genotypes play a role in meiotic and genomic stability in allohexaploids, and highlight the importance of starting with a broad genetic base in establishment of a new allohexaploid *Brassica* crop.

Crosses between *Brassica* allohexaploid types have never before been systematically attempted, and we were curious to see if these behaved as “interspecific” crosses due to the different species parents, or as “intraspecific” crosses due to the shared combination of A, B and C genomes. Pre- and post-fertilization barriers were found to be active in *Brassica* allohexaploid crosses, as many siliques failed to develop and very few seeds were obtained from a number of cross combinations. Many of the plants used in the crosses had viable pollen, but only approximately half of the crosses resulted in developed siliques. In previous studies of diverse crosses between *Brassica* species and ploidy levels, the main barrier to producing new hybrid seeds was the failure of pollen to fertilize the ovule (Nishiyama et al. 1991), and in our crosses, a similar phenomenon might have hindered success in the crosses. The average number of self-seeds produced per silique was higher compared to the values obtained from crossing, also suggesting some incompatibilities when trying to cross between different allohexaploid types with diverse species origin. Out of the five naponigra genotypes, we only obtained self-pollinated seeds from genotype N8.I2. However, we also identified genotypes of *Brassica* naponigra allohexaploids that were more successful at crossing than at producing self-seeds. The differences between self-pollinated and cross-pollinated seed production were especially evident for genotype N8.I3, a genotype that was used as a female and male parent in different crosses but which produced no self-pollinated seeds from three cutting-derived plants (with three selfing bags per plant). In previous studies of the same *Brassica* naponigra genotypes, irregular meiosis was suggested as the main cause of low seed set, while self-incompatibility (present in the *B. nigra* parent species) was thought to have played a smaller role (Gaebelein et al. 2019a). In our study, self-incompatibility seems to have stronger importance, since the same plant was able to produce over 100 seeds from crossing, but under self-pollination conditions not a single seed was developed. We also identified genotypes with significant maternal and paternal influence on number of seeds produced per bud pollination in our crossing scheme. Maternal effects have also been observed in crosses between *B. napus* and *B. rapa,* where the cross has a greater success when *B. napus* is used as a female parent (Fitzjohn et al. 2007). Similarly, when recreating *B. napus* synthetics, maternal *B. rapa* parent genotype strongly influenced the success of the interspecific cross with *B. oleracea* (Diederichsen and Sacristan 1994; Lu et al. 2001; Abel et al. 2005).

Interestingly, 3/7 of our “control” *Brassica* genotypes, representing established allotetraploid species, also contained CNV events. These events were not observed in all five plants analyzed per genotype combination, and some CNVs affected only one of the homologous chromosomes present, suggesting ongoing genomic changes and instability in natural *Brassica* allotetraploids. In support of our results, non-homologous translocations between the A and C genomes have previously been observed in *B. napus* using various methods, including microscopy (Osborn et al. 2003; Sheidai et al. 2003) segregation distortion of mapping populations (Schranz and Osborn 2000; Stein et al. 2017), and sequence read mapping depth (Chalhoub et al. 2014; Samans et al. 2017). Non-homologous pairing in polyploids has also been observed in several other polyploid species, although the frequency is rather low compared to synthetic polyploids (Madlung et al. 2005; Henry et al. 2014; Ihnien Katche et al. 2022) and, even though different chromosomes might interact during the initial stages of meiosis, these configurations tend to resolve into correct homolog pairing (bivalent formation) as meiosis progresses (Comai et al. 2003). How meiotic stabilization or diploid-like behavior has been achieved in natural polyploids is not yet fully understood, although many potential hypothesis have been described, involving for example new mutations, changes in genetic regulation, or inheritance of pre-adapted alleles (reviewed in (Gonzalo 2022)). To the best of our knowledge, our study is the first to report genomic instability in a *B. carinata* genotype: this was unexpected due to the wider genetic divergence between the *Brassica* B and C genomes relative to the A and C genomes (Parkin et al. 1995, Lagercrantz and Lydiate 1996, Perumal et al. 2020). It is possible that this finding relates to the fact that this *B. carinata* genotype is derived from microspore culture to produce doubled-haploid lines, which may confer genomic instability (Shrestha et al. 2023). However, as both *B. carinata* genotypes used in this study are derived from the same microspore culture process (also at the same time point with the same protocol), there may also be variation for non-homologous chromosome recombination frequency within genotypes of the parent allotetraploid species, as has been hypothesized on the basis of cytological results for *B. napus* genotypes (Sheidai et al. 2003). Possibly, such non-homologous rearrangements in the allotetraploid parent species frequently occur but are usually removed due to selection pressure, rather than accumulating in parent lines (Gaeta and Pires 2010).

*Brassica* allohexaploids showed a propensity to reduce genome size. This was observed by the higher number of deletions and missing copy events present in the allohexaploids compared to duplications and extra copies, affecting both whole-chromosomes and chromosome-fragments. Similar trends have also been observed in other allohexaploid studies, such as in carirapa H_2_ lines, and in three NCJ allohexaploid populations, where the majority of the individuals lost chromosomes in subsequent generations (Tian et al. 2010; Gaebelein et al. 2019b). Cultivated *Brassica napus* accessions also showed more deletion events compared to duplication events (Higgins et al. 2018), suggesting that the reduction in genome size is not only an attribute of synthetic material but also may occur in “natural” genotypes. This reduction in genome size could also be an early sign (or ongoing process) of diploidization, similar to what is observed in natural polyploids, where duplicated regions are being lost and rearranged (Li et al. 2021). However, as seen in other studies (Stein et al. 2017; Higgins et al. 2018), duplication/extra copy events are harder to identify using genotyping data, and this might have also influenced our results: hence, more research is needed to see if the propensity to lose chromosomes and chromosome fragments is maintained in future generations.

Accumulation of CNVs from previous generations had a major effect on genome stability: we found different degrees of stability between sibling lines based on the number of accumulated CNVs (Figures 2 and 3). In other words, many lines which originated from the same genotype combination differed from each other in stability and number of accumulated CNVs. This was particularly true for genotype N1C1.J1. Genotype N1C1.J1 contained line 2, which was putatively the most stable line out of the whole set of allohexaploids analyzed: three out of five plants had the expected number of chromosomes (2n = 54), and the other two plants lost only one copy of chromosome A04 or C02, respectively. At the same time, line 2 of N1C1.J1 only had one new CNV event while the remaining events present corresponded to events previously identified in the parent *B. napus* “N1” genotype and hence likely inherited directly from the parent into the allohexaploid. Since the majority of the new CNVs found in N1C1.J1 line 2 appear to be directly inherited from *B. napus,* it is possible that these rearrangements played a beneficial role in stabilizing the karyotype of the new allohexaploids. However, no specific translocation was found that uniquely differentiated “stable” from “unstable” lines within this genotype combination. Interestingly, in synthetic *B. napus* a deleted region in chromosome C01 from 2.5 – 8.3 Mb was previously identified to be associated with lower numbers of seeds produced per flower (Ferreira de Carvalho et al. 2021), although in our case the deleted region in both N1C1.J1 and *B. napus* “N1” was a bit smaller (from 2.2 - 3.3 Mb on chromosome C01), and did not seem to affect seed production.

An overall bias in translocation direction between the A and C subgenomes was observed for fixed non-reciprocal translocation events, where significantly more instances were found where a C-genome fragment replaced an A-genome fragment, compared to where an A-genome fragment replaced a C-genome fragment. However, at the genotype level, there were different patterns of bias in either A → C or C → A translocations, with differences also found between lines within genotypes. For example, genotype N1C1.J1 (containing the putatively stable line 2) had only fixed translocation events where an A fragment replaced a C fragment, while genotype N5C2.J2a had more fixed events where a C fragment replaced an A fragment as a result of non-homologous recombination. In natural and synthetic *Brassica napus* an opposite trend has been observed, with the A genome replacing the C genome more frequently than the C genome replaced the A (Samans et al. 2017; Higgins et al. 2018). Interestingly, translocations in allohexaploids where an A fragment replaced a C fragment have previously been associated with a positive effect on fertility (Gaebelein et al. 2019b); this effect was previously found for genotype N5C2.J2. For this same genotype in our study (N5C2.J2a), we observed the opposite of this translocation bias effect to the later generations (more C fragments translocated into A chromosomes), although we selected for high-fertility genotypes in our study.

Four different fixed putative non-reciprocal translocations between a B-genome chromosome and a C-genome chromosome were found, B genome introgressions obtained in *B. napus* were also found to be more common in the C genome compared to the A genome (Dhaliwal et al. 2017); in this study, 17 out of 23 B-genome segments identified were introgressed into the C subgenome: eight involved chromosomes B06 or B07, while the remainder could not be identified to the chromosome level. In interspecific hybrids between *B. napus* × *B. carinata* followed by two rounds of backcrossing, there was only one indication of a B-chromosome segment introgression involving a portion of chromosome B05 which introgressed into either chromosome A01 or C01, while the remaining B chromosomes were either missing or present as whole additional chromosomes (Navabi et al. 2011). In our study, only one event involving translocation of an A genome fragment into the B subgenome was found. However, care should be taken about generalizations related to frequency of C vs. A-genome introgressions of B-genome segments, due to the low numbers of these events observed in our study.

We also observed the apparent loss of B-chromosome fragments or whole chromosomes without evidence for an accompanying duplication/translocation event. B genome chromosome loss was primarily observed for three chromosomes: B04 (NCJ genotypes), B05 (NCJ and carirapa genotypes), and B07 (NCJ genotype). NCJ allohexaploid types were produced via a two-step crossing process, whereby some loss of univalent A- and B-genome chromosomes from the *B. napus* × *B. carinata* (CCAB hybrid) likely occurred (Mason et al. 2010, 2012) . However, whether there is specific selection against these three B chromosomes in particular (especially B07, which was also lost in carirapa lines) is unknown. Navabi et al. (2011) also found loss of whole B chromosomes and deletions in terminal regions (Navabi et al. 2011), which may suggest preferential loss of these chromosomes and translocation regions, although more data is needed to confirm this result.

Despite intensive selection for several generations, most of the allohexaploid plants analyzed were aneuploids, and we were only able to identify seven plants with the expected number of chromosomes. Most likely, this can be attributed to the means of selection in each generation, which was fertility (number of seeds produced) rather than euploidy. Although fertility is known to be correlated with regularity of meiotic behavior in allohexaploids (Gaebelein et al. 2019b), many allohexaploids have also been observed to show high fertility despite presence of large number of CNVs (Mason et al. 2014, 2015). In previous studies in *B. napus* synthetics, the number of euploid plants increased per generation upon successive selection of euploid individuals for several generations, with initial generations S_1_-S_3_ having 88.5 - 88.7% euploid plants, and later generations (S_8_) having 100% euploid plants (Ferreira de Carvalho et al. 2021). *Brassica napus* synthetics that are not under generational selective pressure show no evidence of karyotype stabilization, and tend to accumulate extra chromosomes (Xiong et al. 2011). We did not observe accumulation of extra chromosomes in the allohexaploid lines, but it might be possible that we underestimated the number of duplication events using our SNP genotyping method. In another study of a subset of wheat synthetic hexaploid lines, aneuploidy was observed as a characteristic of synthetic nascent hexaploid lines and, despite selection for euploidy in early and subsequent generations, karyotype stabilization and reduction of aneuploidy was not achieved (Zhang et al. 2013). In *Tragopogon* allopolyploids, aneuploidy is also still frequently observed after approximately 40 generations (Chester et al. 2012, 2015), suggesting pressure to stabilize meiosis may also not always be present in natural systems. In early generations (H_2_) a large set of *Brassica* carirapa allohexaploids produced from crossing several different accessions (29 *B. rapa* and 107 *B. carinata*) showed high levels of aneuploidy (91%) and low levels of putative euploid plants (4.6%) (Tian et al. 2010). Although euploidy was rare in our study, it is possible that more stringent selection in each generation for euploid chromosome complements might improve generational stability.

Viviparous seeds were observed in most of the crosses we made. We also identified significant differences in the number of viviparous seeds obtained based on the maternal genotype used in the cross. Similarly, in previous studies of crosses involving *B. napus* and *B. rapa,* viviparous seeds were also obtained (4.2%), but only when *B. rapa* was used as the female parent (Hauser and Østergåurd 2004; Jenkins et al. 2005). In previous studies it was proposed that the germination of seeds inside the silique could be a response to incompatibility between the new hybrid seed (maternal and paternal genomes) and its new silique environment (determined by the maternal genome) (Hauser and Østergåurd 2004). Although the exact reason why viviparous seeds occurred so commonly in our crossings requires further study, it is important to account for a potential loss of hybrid seeds due to this premature germination, which also makes the resulting seeds more prone to drought, fungal infection or simply death (Jenkins et al. 2005).

Out of the eight new selected F_1_ hybrids, only the hybrid C13b × C05 produced significantly more seeds (heterosis) than both parents, possibly because C13 and C05 were both originally 100% homozygous genotypes (in contrast to NCJ types, which are heterozygous in the first generation). In the case of the hybrid N6C2.J2a × J3O1a, despite having no significant differences in total number of CNV between parents and hybrids, the parent N6C2.J2a produced significantly more seeds than parent J3O1a and the respective hybrid. We also observed that in a cross between N8.I3 × J3O1, the F_1_ hybrids had significantly lower total numbers of CNVs compared to the parent J3O1. In previous studies of F_1_ hybrids between carirapa and NCJ type allohexaploids, new F_1_ hybrids accumulated and produced more novel rearrangements compared to the parents despite potential disadvantages in meiosis (Quezada-Martinez et al. 2022). The lack of heterosis in our F_1_ hybrid study is not surprising, since it was been observed that the main factor affecting fertility (seed number) in *Brassica* allohexaploids is the presence of absence of rearrangements (Gaebelein et al. 2019b).

## Conclusions

Our results suggest that genomically stable synthetic *Brassica* allohexaploids are achievable, but only at extremely low frequencies, and that stability may not always be under positive selective pressure due to the unpredictable relationship between fertility and genome composition in these hybrid types. Combining different allohexaploid types via hand-pollination is feasible, allowing new allelic combinations to be produced. However, the majority of the F_1_ hybrids analyzed in the present study showed no significant improvement in fertility or genome stability compared to their parents. Translocations between the A-B and C-B subgenomes were also observed, and may be valuable for use in introgression breeding programs.

## Author Contribution Statement

DQM carried out the experiments, collected and analysed the data, produced the figures and tables and drafted the manuscript. JB generated data and contributed to critical revisions of the manuscript. ASM acquired funding, conceptualized the project with DQM, supervised DQM and contributed to critical revisions of the manuscript.

## Data Availability Statement

All data produced as part of this project is available in the supplementary files.

## Acknowledgements

We gratefully acknowledge Ingrid Schneider-Hüther and Liane Renno for technical assistance and plant management. We also thank Poonam Bangia for her contribution in seed harvesting.

## Supplementary Figures

**Supplementary Figure 1.** Allohexaploid lines ranked from low to high total number of single-copy events affecting chromosome counts. *Brassica* allohexaploid genotypes NCJ: N1C1.J1, N1C2.J1, N5C2.J2a, N5C2.J2b; Carirapa: C05, C13, C2128, and junleracea: J3O1. Each line per genotype is shown as “_” followed by the number of the line (1 to 5). Each line represents the total number of events in all the individuals analyzed (located in the centromere region, extra copy or missing copy).

**Supplementary Figure 2:** Frequency of copy number variation (CNV) event size per allohexaploid type. Allohexaploid type is shown according to the legend: NCJ in pink, carirapa in purple, and junleracea in dark green. The size is represented in megabases (Mb) distributed in 1Mb bin width for each of the CNV type: deletion, duplication, missing copy, and extra copy.

**Supplementary Figure 3.** Size comparison between allohexaploid types for different copy number variation (CNV) events. Each dot represents a single event scored per allohexaploid type. Allohexaploid type is colored according to the legend: NCJ in pink, carirapa in purple, and junleracea in dark green. Mean values per group is shown by a red horizontal line. Comparisons were made within the groups and between the allohexaploid type (Kruskal-Wallis test: deletion *p* = 1.3e-19, duplication *p* = 1.4e-6, missing copy *p* = 1.3e-6, extra copy *p* = 4.7e-3). Multiple pairwise comparisons were done using Dunńs test with the level of significance shown in brackets as: “ns” (not significant) = *p* > 0.05, “*” = *p* ≤ 0.05, “**” = *p* ≤ 0.01, “****” = *p* ≤ 0.0001.

**Supplementary Figure 4.** Percentage of pollen viability in the different allohexaploid genotypes used for crossing. Each dot represents the percentage of pollen viability per individual from the *Brassica* allohexaploid genotypes NCJ: N1C1.J1, N1C2.J1, N5C2.J1, N5C2.J2a, N5C2.J2b, N6C2.J2, N7C1.J1; Carirapa: C05, C13, C2128, naponigra: N8.I2, N8.I3, N5.I2; and junleracea: J3O1. Mean per genotype is represented by a red triangle.

**Supplementary Figure 5:** Total seeds produced by parental genotypes (P1 and P2) and their corresponding F_1_ hybrid in *Brassica* allohexaploids. The genotype combination from each of the F1 hybrids is represented at the top of each graph. P1 and P2 correspond to the genotype used in the combination and are the first and second genotype named in the cross, respectively. Each dot represents one plant per group. Statistical differences (Student’s pairwise t-test, unpaired t-test) between parents and hybrids are represented by brackets. Ns = Not significant. NA = Not analyzed. Average value per group is represented by a red horizontal line. NA = genotype not analyzed.

**Supplementary Figure 6:** Total copy number variants (CNVs) in allohexaploid parents compared to their F_1_ hybrids. Each dot represents one plant per group. Statistical differences between parents and hybrids are represented by brackets (Dunńs test), ns = not significant. NA = not analyzed. Average value per group is represented by a red horizontal line.

## Supplementary Tables

**Supplementary Table 1:** Plant material used for genotyping in the genome stability analysis. Allohexaploid types are the product of crossing different parental species: NCJ = *B. napus* × *B. carinata* × *B. juncea;* carirapa = *B. carinata* × *B. rapa;* and junleracea = *B. juncea* × *B. oleracea.* The particular parental species, genotype, and code used for publication is described in the table. Genotype N5C2.J2 has two different lineages, represented by “a” and “b” following the genotype name.

**Supplementary Table 2:** Plant material used for the crossing of different allohexaploid genotypes. Allohexaploid types are the product of crossing different parental species: NCJ = *B. napus* × *B. carinata* × *B. juncea;* carirapa = *B. carinata* × *B. rapa;* naponigra = *B. napus* × *B. nigra*, and junleracea = *B. juncea* × *B. oleracea.* The particular parental species, genotype, and code used for publication is described in the table. Genotype N5C2.J2 has two different lineages, represented by “a” and “b” following the genotype name. Days to flower were counted from the day of sowing until first flower open (BBCH60). Pollen viability was measured using two different methods: Amphasys or acetocarmine staining (*). Self-seeds were obtained by bagging 1-10 branches per plant. Ratio was calculated by dividing self-seeds obtained by the number of siliques harvested.

**Supplementary Table 3:** Crossing scheme planned to combine different allohexaploid genotypes. Parent 1and 2 columns are defining the genotypes intended to be cross but not the direction of the cross. Allohexaploid types are the product of crossing different parental species: NCJ = *B*. *napus* × *B. carinata* × *B. juncea;* carirapa = *B. carinata* × *B. rapa;* naponigra = *B. napus* × *B. nigra*, and junleracea = *B. juncea* × *B. oleracea*.

**Supplementary Table 4:** Genotype and Log R ratio data for *Brassica* allohexaploid genotypes used in the genome stability and F1 hybrid analysis. Genotype data obtained from the 90K *Brassica* SNP chip, with markers distributed in chromosomes from the subgenomes A (A01-A10), C (C01-C09), and B (B01-B08). Biological replicates have the same genotype name.

**Supplementary Table 5:** Outcome of the linear regression model analysis to estimate (a) genotype specific effect, (b) allohexaploid type, (c) *Brassica napus* genotype effect, (d) *Brassica carinata* genotype effect, (e) *Brassica juncea* genotype effect in the total of CNVs in NCJ allohexaploids.

**Supplementary Table 6:** Outcome of the multiple linear regression model analysis to estimate the effects of maternal (♀) and paternal (♂) genotype used in the crossing of different *Brassica* allohexaploids. Allohexaploid types are the product of crossing different parental species: NCJ = *B. napus* × *B. carinata* × *B. juncea;* carirapa = *B. carinata* × *B. rapa;* naponigra = *B. napus* × *B. nigra*, and junleracea = *B. juncea* × *B. oleracea.* In green are highlighted the genotypes with a significant effect.

**Supplementary Table 7:** Outcome of the multiple linear regression model analysis to estimate the effect of *Brassica* parents in common in a cross, together with the maternal (♀) and paternal (♂) genotype effect within NCJ allohexaploid genotypes. NCJ allohexaploids were produced by the cross between *B. napus* × *B. carinata × B. juncea.* In green are highlighted the genotypes with a significant effect.

**Supplementary Table 8:** Outcome of the linear regression model analysis to estimate the maternal parent effect in the production of viviparous seeds after crossing different *Brassica* allohexaploids. NCJ allohexaploids were produced by the cross between *B. napus* × *B. carinata × B. juncea.* Highlighted in yellow are the genotypes with a significant effect.

**Supplementary Table 9:** Crosses between *Brassica* hexaploids NCJ (*B. napus* × *B. carinata* × *B. juncea*), junleracea (*B. juncea* × *B. oleracea*), carirapa (*B. carinata* × *B. rapa*), and naponigra (*B. napus* × *B. nigra*) genotypes selected for comparison with the parents. The individual parental plant used in the cross is represented by a hyphen followed by the number of the plant (*). Self-seeds were produced from the total plant. Days to flower were calculated from the day of sowing until the opening of the first flower (BBCH60).

